# Three dimensional reconstruction of the human nucleus accumbens reveals topographic organization and molecular heterogeneity of D1-islands across the anterior posterior axis

**DOI:** 10.64898/2026.09.25.754413

**Authors:** Robert A. Phillips, Jianing Yao, Svitlana V. Bach, Ishbel Del Rosario Alvia, Yufeng Du, Sarah E. Maguire, Ruth Zhang, Ryan A. Miller, Joel E. Kleinman, Thomas M. Hyde, Keri Martinowich, Kristen R. Maynard, Stephanie C. Hicks

## Abstract

Spatial transcriptomics has transformed molecular characterization of the human brain, but most studies profile individual two-dimensional tissue sections and therefore cannot capture molecular organization across 3D neuroanatomical axes. Here, we developed a spatial genomics framework to reconstruct the 3D cellular architecture of the human nucleus accumbens (NAc) by densely sampling and aligning serial sections across its anterior-posterior (AP) extent. Combining single-cell Xenium profiling with near-transcriptome-wide VisiumHD, we mapped cell type composition and topography across AP, mediolateral and dorsoventral axes. Neuronal populations varied in relative abundance across the AP axis while maintaining characteristic spatial topographies. These differences were particularly pronounced among D1-islands, a topographically organized structure within the NAc in which transcriptionally distinct subpopulations of *DRD1*-expressing medium spiny neurons (MSNs) aggregate into discrete cellular formations. D1-islands comprised two molecularly distinct spatial domains with conserved counterparts across species. Both domains varied in relative abundance across the AP axis, but demonstrated distinct 3D distribution; one was present throughout the NAc, whereas the other, corresponding to islands of Calleja (ICj), was more restricted to intermediate-to-posterior levels. Local spatial analyses revealed region- and AP-dependent cellular relationships, including segregation of the two island populations despite their anatomical proximity. Together, these findings establish a framework for three dimensional spatial genomics of human brain tissue and reveal molecular and cellular organization not captured by individual tissue sections.

## 1 Introduction

The nucleus accumbens (NAc) is a ventral striatal structure that integrates dopaminergic input from the ventral tegmental area (VTA) to regulate reward, motivation, and goal-directed behavior. Medium spiny neurons (MSNs), the principal neuronal population of the NAc, have traditionally been classified based on expression of D1 or D2 dopamine (DA) receptors. However, single-cell and spatial transcriptomic studies have revealed substantial molecular heterogeneity within these canonical MSN classes^1–8^, and demonstrated that molecularly distinct subtypes exhibit characteristic spatial topographies across the NAc^1,2,6,8^. Rodent circuit-mapping studies have further shown that afferent connectivity can influence topographic organization of MSNs, underscoring potential relationships between spatial organization and circuit architecture^9,10^.

A striking example of this spatiomolecular organization is a subpopulation of molecularly distinct D1-MSNs, termed D1-islands, that form dense clusters along the medial and ventral borders of the NAc. D1-island populations have been identified across rodents^1,6,11^, non-human primates (NHP)^8^, and humans ^2^, where they share characteristic spatial organization and conserved molecular features, including prominent expression of the mu-opioid receptor gene, *OPRM1*. In rats, these cells have been described as Chst9-MSNs^1,3,4,11^, which in addition to *Oprm1*, express high levels of *Chst9* and *Grm8*. Within the rat, a separate population of D1-islands exist and are termed Sema5a-MSNs due to their high expression of *Sema5a*^1^. Chst9- and Sema5a-MSNs are molecularly distinct and are differentially enriched across the AP and ML axes^1^. A related population in NHPs, termed D1-neurochemically unique domains in accumbens and putamen (D1-NUDAP)^8^, is characterized by high expression of *RXPF1, CPNE4* and other conserved markers such as *OPRM1*. In the human NAc, we identified similarly organized D1-islands enriched for *FOXP2, RXFP1*, and *GABRQ*^*2*^. Beyond their conserved molecular and spatial organization, rodent studies have demonstrated that molecularly distinct MSN cells within D1-islands regulate local DA release^12^ and contribute to opioid-dependent reward learning and withdrawal-related behaviors^1,12^. Importantly, human D1-islands are enriched for genes associated with genetic risk for bipolar disorder, schizophrenia and major depressive disorder ^2^, further supporting the potential relevance of this population in human brain function and disease.

Despite growing recognition of the molecular, cellular, and spatial heterogeneity in the NAc, most studies have investigated the topography of NAc cell types at a single or limited number of two-dimensional tissue sections. In our initial spot-based spatial transcriptomics study profiling the human NAc, we showed that D1-islands differed in both location and abundance across tissue sections sampled at different anterior-posterior (AP) positions, providing potential evidence for heterogeneity across this anatomical axis in human brain^2^. However, AP positions were not systematically sampled within the same brain across individuals, limiting our ability to distinguish AP-dependent organization from inter-individual variability. Understanding this topographic organization is important because anatomical position within the NAc is closely related to circuit organization and behavioral function. Afferent projections to the NAc exhibit distinct topographic organization^13–15^, while various molecularly-defined neuronal populations have been linked to distinct reward or aversion-related behaviors^16^. Similarly, opioid-sensitive “hotspots” localize to discrete anatomical regions within the ventral striatum, where opioid signaling produces location-dependent effects on hedonic processing^17^. Thus, understanding where molecularly defined populations are positioned within the human NAc may provide important insights into the circuits in which they participate and their potential functions. Comprehensive profiling across the AP axis is therefore needed to determine how the abundance, topography and local cellular environments of D1-islands and other molecularly distinct cell populations vary across the NAc’s 3D structure.

Here, we used Xenium spatial transcriptomics to systematically profile the molecular and spatial organization of cell types within the human NAc. We profiled 22 tissue sections from 2 neurotypical donors sampled at approximately 500 µm intervals across the full AP extent and developed a computational framework to reconstruct cell type organization in three dimensions (3D). We complemented this approach with targeted VisiumHD profiling to obtain near-transcriptome-wide molecular characterization of spatially defined D1-island populations. These analyses resolved two molecularly distinct island populations with different 3D distributions, including an *OPRM1*/*RXFP1*/*GABRQ*-expressing population corresponding to previously described D1-islands, and a *PROK2*/*VIP*-expressing population that we identified as human Islands of Calleja (ICj). Cross-species analyses further demonstrated molecular conservation of both D1-island and ICj populations across rodents, NHPs and humans. Together, this study provides a 3D spatiomolecular map of the human NAc and demonstrates that molecularly distinct NAc populations exhibit differences in abundance and local cellular organization across its AP extent.

## 2 Results

### 2.1 Identification of topographically organized, transcriptionally distinct cell types across the anterior-posterior (AP) axis of the human NAc

We previously identified that transcriptionally and spatially distinct cell types are organized in gradients across the human NAc’s medial-lateral (ML) and dorsal-ventral (DV) axes^2^, and provided initial evidence for changes in cell type composition across the anterior-posterior (AP) axis. However, in that study the NAc was sampled at different AP positions across the 10 donors, confounding anatomical location with samples. Hence, it remained unclear how transcriptionally distinct cell types, especially those forming D1-islands, differed along the AP axis^2^. To systematically profile across the AP axis, we first identified two neurotypical control donors where the entire NAc was retained within a single coronal fresh-frozen brain slab (**Fig. S1, Fig. S2**). We then performed Xenium *in situ* sequencing (10x Genomics) on tissue sections collected at 500 µm intervals along the entire AP extent (**Fig. 1A**; n=11 sections/donor). Xenium is an imaging-based spatially-resolved transcriptomics (SRT) technology that uses a probe hybridization strategy for visualizing targeted genes at single cell resolution. Using data from our earlier transcriptomics studies of the human NAc^2,5^, we developed a 100 gene custom panel combined with a 266 gene “human brain” base panel to target 366 total genes (**Table S1**). The custom panel contained top marker genes for neuronal subtypes with 12 genes specific to previously identified D1-island subtypes (**Table S1**). Other targeted genes included those marking surrounding structures (e.g. lateral septum and cortex), spatially variable genes specific to NAc spatial gradients previously described by our group^2^, as well as risk genes for schizophrenia and opioid use disorder (OUD). Following quality control measures (**Fig. S3**), non-spatial clustering identified 20 transcriptionally distinct cell types (**Fig. 1B**), including neuronal and non-neuronal subtypes that occupy distinct topographical domains along the ML/DV axes (**Fig. 1B-D**). We identified two populations of *DRD1*-expressing MSNs (D1_Island_A and D1_Island_B), which formed tight clusters across the AP axis with unique topographical patterns along the dorsomedial, ventral-medial, and ventral-lateral borders, and exhibited transcriptional signatures distinguished by unique receptor and peptide expression profiles (**Fig. 1B,E**). D1_Island_A selectively expressed *RXFP1*, which encodes relaxin peptide 1, and *TRHDE*, which encodes thyroid hormone degrading enzyme (**Fig. 1E**). D1_Island_B selectively expresses *PROK2*, which encodes the secreted protein prokineticin-2, and *VIP*, which encodes vasoactive intestinal peptide (**Fig. 1E**). While both D1-island cell types expressed *TZHZ1* and *CPNE4*, which mark D1-islands in rodents and non-human primates (NHPs), expression was higher in D1_Island_A compared to D1_Island_B (**Fig. 1E**). Both Xenium D1-island subtypes correlated with the D1-island spatial domain identified in our spot-based Visium SRT dataset of the human NAc^2^ (**Fig. S4**), suggesting that this previously identified spatial domain likely contained two transcriptionally distinct cell types which could only be resolved with cellular resolution. In addition to identifying D1-island subtypes, we also defined *DRD1*-expressing MSNs (DRD1_MSN) based on their expression of *DRD1*/*TAC1*/*RELN*/*PDYN* and *DRD2*-expressing MSNs (DRD2_MSN) based on their expression of *DRD2*/*ADORA2A*/*PENK*/*GPR6*. Correlation of transcriptional signatures between Xenium cell types and previously published MSN subtypes ^2^ found that DRD1_MSN and DRD2_MSN likely contained several populations of DRD1- and DRD2-MSN subtypes (**Fig. S4**). This interpretation is further supported by the fact that DRD1_MSN and DRD2_MSN did not occupy a specific topographic domain, but rather were evenly distributed across the DV, ML, and AP axes of the human NAc (**Fig. 1B**). In addition to MSN subtypes, we also identified canonical GABAergic neuron subtypes (Inh_PVALB and Inh_SST), as well as cholinergic neurons expressing *CHAT*, which were located within the NAc and neighboring regions (**Fig. 1B**). Additionally, we found *SLC17A7-*expressing excitatory neurons ventral to the NAc in adjacent cortical regions (**Fig. 1B,E**). While our custom gene panel was primarily designed to resolve neuronal heterogeneity, we also identified several non-neuronal populations including astrocytes and ependymal cells (**Fig. 1E**). Specifically, we identified two populations of astrocytes (Astro_A and Astro_B) marked by *GJA1* and *AQP4*, which differed based on expression of *GFAP, TNC*, and *WIF1*. We also identified two populations of Fibroblasts (Fibro_A and Fibro_B) that differed based on expression of *CLDN5* and *NR2F2* (**Fig. 1E**). Three of the distinct cell types identified by non-spatial clustering represented mixtures of MSNs, astrocytes, and microglia (**Fig. 1E**). The mixed cellular composition of these clusters likely reflected limitations in cell segmentation and transcript quantification.

**Figure 1.**
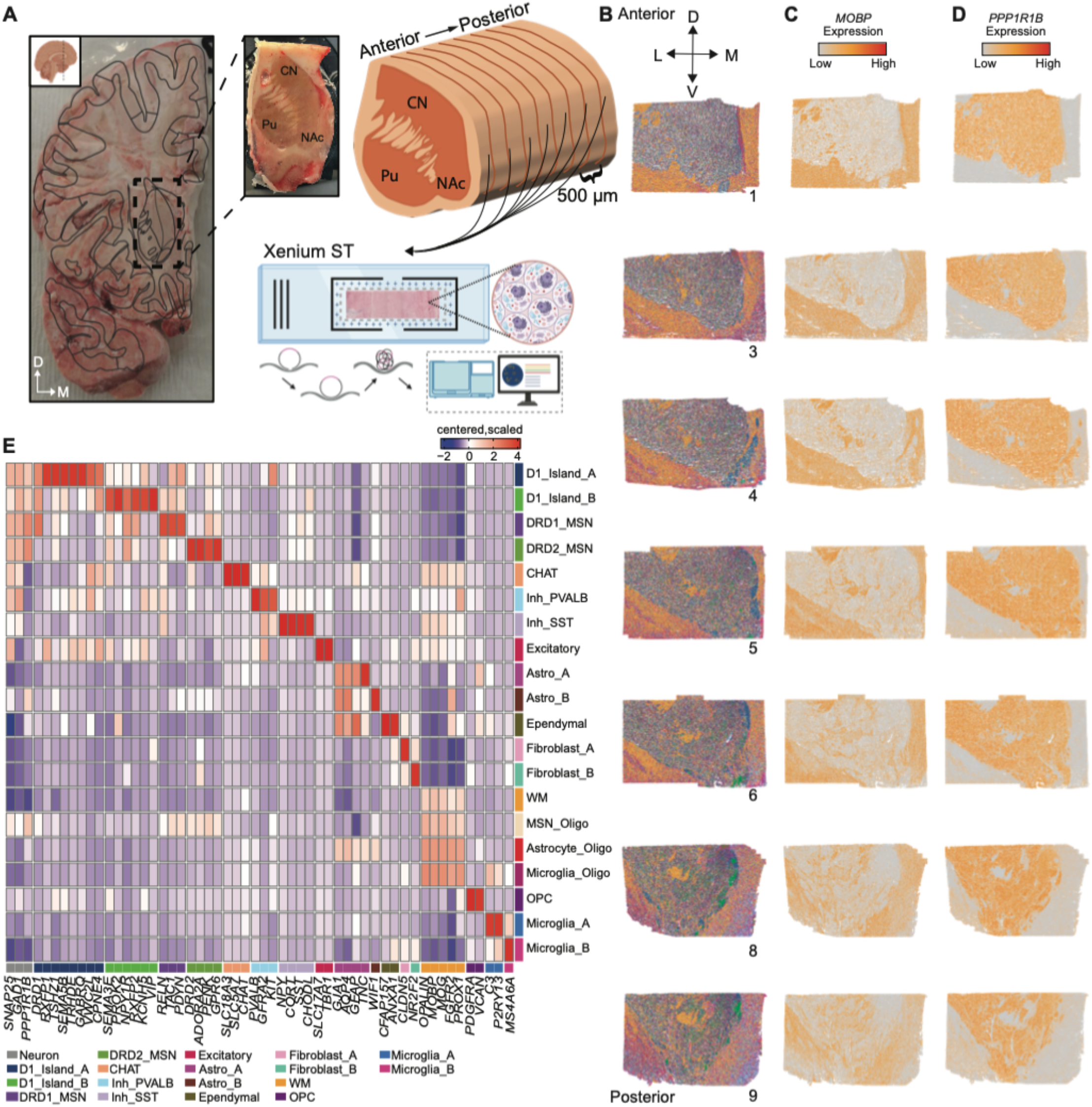
Identification of transcriptionally distinct cellular populations across the AP axis of the human NAc. **A**. Overview of experimental design. Cryosections were collected at 500 µm intervals across the AP extent of the NAc from two neurotypical control brain donors (1M/1F) onto Xenium slides and probed with a 366 gene panel. **B**. Tissue sections colored by 20 molecularly distinct cell types derived from non-spatial *BANKSY*^*60*^ clustering. The number at bottom right of each section indicates slice number out of 11 total. **C**. Tissue sections with cells colored by expression of *MOBP*, marking white matter. **D**. Tissue sections with cells colored by *PPP1R1B* expression, a marker of medium spiny neurons (MSNs). **E**. Heatmap of expression profiles of *BANKSY* cell types.

### 2.2 Molecularly distinct populations of D1-islands exhibit dynamic changes in cell type proportion across the AP axis

Consistent with our previous work2, the abundance of some cell types appeared to vary along the AP axis of the NAc, most notably the two D1-island populations (**Fig. 2A**). Examining their three-dimensional organization requires reliable anatomical correspondence across consecutive tissue sections. However, tissue banking procedures can complicate this correspondence: rapid slabbing of fresh tissue can produce slabs that are not perpendicular to the AP axis, and freezing can introduce anatomical distortions. Investigation of the tissue block and surrounding anatomical landmarks revealed that the assayed sections from the male donor were cut from an angled slab. This complicated anatomical interpretation across AP levels and obscured precise NAc borders in posterior sections. We therefore restricted the following 3D analyses to the female donor, Br6660, whose sections were cut approximately in the coronal plane and had well-preserved anatomy. To quantify the topographic organization of cell types across the NAc AP axis, we generated a 3D reconstruction by aligning 11 consecutive Xenium tissue sections from this donor using *Spateo*^18^ (**Fig. 2B, Fig. S5, Video S1**).

**Figure 2.**
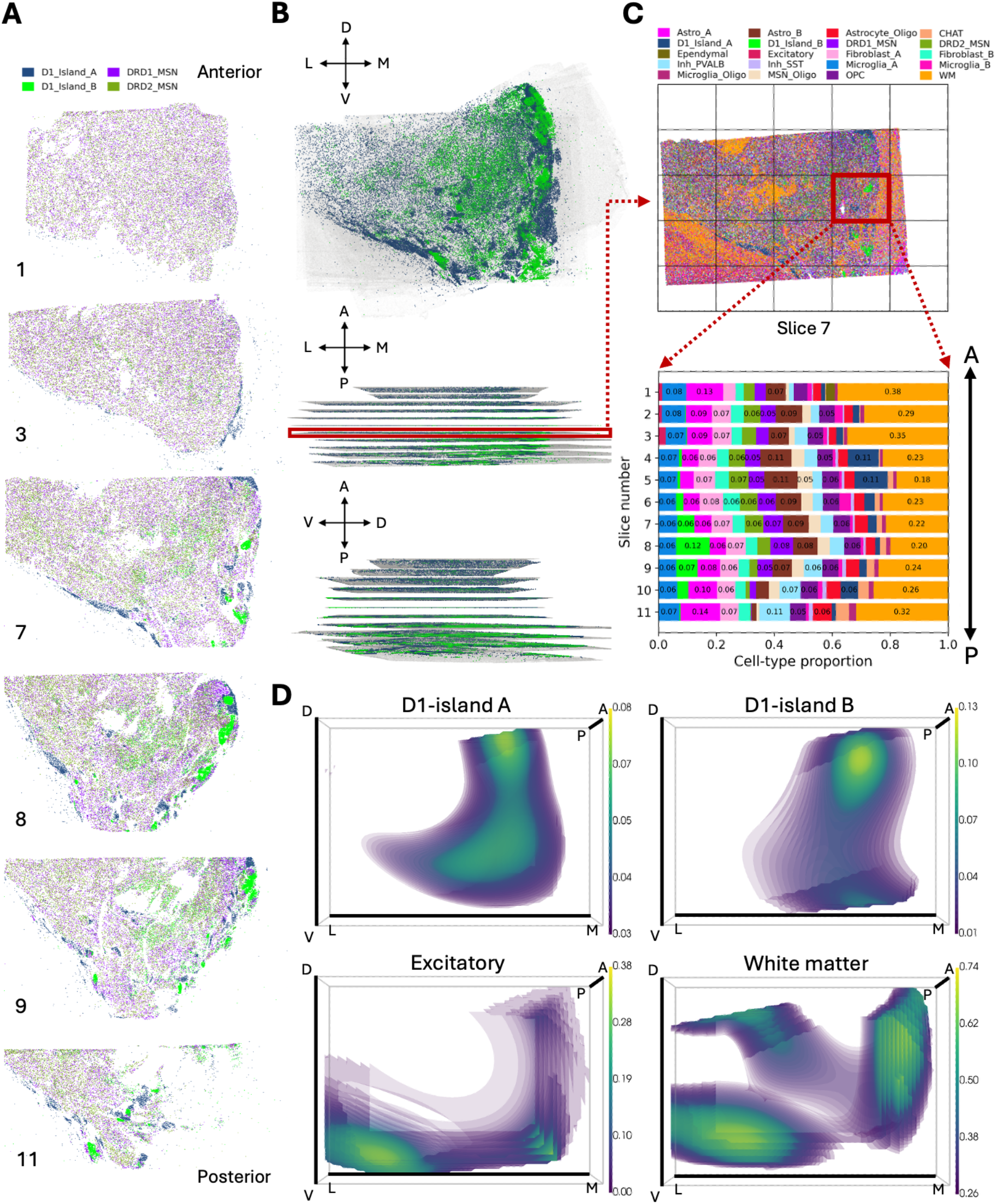
Cell type composition and spatial organization vary along the anterior–posterior axis. **A**. Spatial distribution of D1-island and MSN populations in representative Xenium tissue slices from donor Br6660, shown from anterior to posterior. The number at bottom left of each section indicates slice number out of 11 total. **B**. 3D reconstruction of aligned Xenium tissue slices from donor Br6660 (*n*=11). The tissue volume is shown from three orientations along the dorsal–ventral (DV), medial-lateral (ML), and anterior–posterior (AP) axes. **C**. A representative tissue slice at an AP depth of 3,580 µm was divided into a 5 × 5 grid comprising 25 tiles. The stacked bar plot shows cell type proportions within the selected medial tile (red box in B) across AP levels. **D**. 3D smoothed surfaces of cell type proportions across the reconstructed tissue volume, estimated using binomial generalized additive models. For each cell type, only locations with fitted proportions above the 60th percentile within the tissue are rendered; lighter colors indicate higher fitted proportions.

We first quantified how cell-type composition varied along the AP axis within corresponding spatial regions. A common 5 × 5 in-plane grid was applied to the aligned sections, and cell-type proportions were calculated within each of the 25 tiles at each AP level (**Fig. 2C, Fig. S6A**). Here, we focused on medial tiles as D1-islands were predominantly located along the medial NAc border (**Fig. S6A**). D1_Island_A was observed throughout the sampled AP axis, whereas D1_Island_B contributed a larger proportion at intermediate-to-posterior levels (**Fig. 2C, Fig. S6B**). In contrast, many other cell types exhibited stable cell type proportions along the AP axis, including GABAergic paravalbumin neurons and oligodendrocytes (**Fig. 2C**). Despite these differences in proportions, the broad topographic organization of D1-island populations remained relatively consistent across sections. For example, D1_Island_B remained predominantly localized near the NAc border across sections in which it was observed, with some variation in its precise position within the ML–DV plane (**Fig. 2A, Fig. S6A**).

Although the grid analysis enabled direct comparisons of cell-type composition across AP levels, summarizing each cell type within a tile by a single proportion did not capture within-tile spatial variation. To visualize this variation and examine how the patterns were organized jointly across the ML, DV, and AP axes, we fitted a binomial generalized additive model for each cell type to estimate local cell-type proportions across the reconstructed tissue volume. The resulting graphs provided an integrated view of the location and spatial extent of regions with higher fitted proportions for each cell type of interest, with lighter colors indicating higher fitted proportions (**Fig. 2D**). At anterior-to-intermediate AP levels, D1_Island_A extended across the ventromedial-to-ventrolateral NAc, whereas at intermediate-to-posterior levels, it was most prominent in a compact dorsomedial domain (**Video S1, Video S2**). We found that D1_Island_B was more restricted to intermediate-to-posterior AP levels and similarly formed regions of higher proportions along the dorsomedial border of the NAc (**Video S1**,**Video S3**). Together with the observed cell distributions, these localized domains were consistent with the spatially discrete, island-like organization identified in our previous spot-based spatial transcriptomics study2. Although D1_Island_A and D1_Island_B occupied partially overlapping anatomical regions, their smoothed surfaces revealed distinct 3D spatial patterns. White-matter, consisting primarily of oligodendrocytes, surrounded the NAc and also formed a tract traversing the reconstructed volume, anatomically consistent with the internal capsule (**Fig. 2D, Video S4**). Excitatory neurons were concentrated immediately ventral to the NAc, forming another spatially distinct domain consistent with adjacent cortical regions (**Fig. 2D, Video S5**). Given that dynamic changes in cell type proportion across the AP axis may be mirrored by transcriptional changes, we next sought to understand AP-associated changes in gene expression. However, none of the 14 D1_Island_A or D1_Island_B marker genes (**Fig. 1E**) showed a statistically significant association with AP level after Bonferroni correction (minimum adjusted *p* = 0.094; **Fig. S7**).

### 2.3 Targeted spatiomolecular profiling of D1-island populations with VisiumHD identifies human Islands of Calleja (ICj)

Given the dynamic cell type proportions of NAc cell types, particularly D1-islands, along the AP axis, we generated more comprehensive transcriptomes of these cell types. Specifically, we used Visium High Definition (VisiumHD), a subcellular sequencing-based SRT technology measuring over 18,000 protein-coding genes, to define transcriptome-wide profiles. Given the unique transcriptional and topographic organization of D1-islands, we specifically targeted 8 regions of interest (ROIs) across the NAc that contained high proportions of these cell types (**Fig. 3A**). Following quality control (**Fig. S8**), segmented cells contained an average of ~813 genes and 1,113 UMIs. We performed spatial clustering using *BANSKY* to identify whether D1-island cell types map to unique spatial domains. We identified 12 spatial domains which included white matter, ventricle-lining ependymal cells, and broad MSN-rich neuronal domains (MSN_Inh_A-C, **Fig. 3A)**. While non-neuronal domains were marked by distinct gene expression signatures, MSN_Inh_A-C displayed an overlapping transcriptional signature characterized by *PPP1R1B*/*DRD1*/*DRD2* as well as *NPY*/*CORT*/*SST* (**Fig. 3C**), indicating that these domains contain both MSNs and inhibitory neuronal subtypes. This was further supported by spatial registration between VisiumHD spatial domains (**Fig. S9**) and previously reported snRNA-seq cell types/Visium spatial domains ^2^, which showed that MSN_Inh_A-C spatial domains were highly correlated with both MSN and inhibitory neuron subtypes (**Fig. S9**). Similar to Xenium SRT, we identified two D1-island spatial domains that consisted of molecularly distinct D1-MSNs which formed tight bundles of cells along the NAc border (**Fig. 3A**). Both spatial domains were enriched for *DRD1*, but differed on several key gene classes. First, D1_Island_A was enriched for *PPP1R1B* and *BCL11B*, two canonical MSN markers (**Fig. 3C**). D1_Island_B was depleted for *PPP1R1B* compared to D1_Island_A, but exhibited high expression of *BCL11B* and *ISL1. ISL1* is a transcription factor required for MSN differentiation, suggesting that D1_Island_B may represent a group of immature MSNs that are similar to the Islands of Calleja (ICj), a distinct subset of neurons that are typically located ventral to the NAc in the olfactory tubercle (OT)^19–21^. The classification of D1_Island_B as ICjJ was further supported by the dense expression of *DRD3*, which has been reported to mark ICj^22–24^. Unexpectedly, we found ICj cells located within the NAc proper, as well as along the medial border between the NAc and surrounding white matter tracts. D1_Island_A also exhibited high expression of *RXFP1, TSHZ1*, and *OPRM1*, genes that mark D1-islands in rodent NAc^1,12^. The enrichment of these genes suggested that D1_Island_A represents a canonical D1-island within the NAc that is made up of DRD1-expressing MSNs, while D1_Island_B represents ICj.

**Figure 3.**
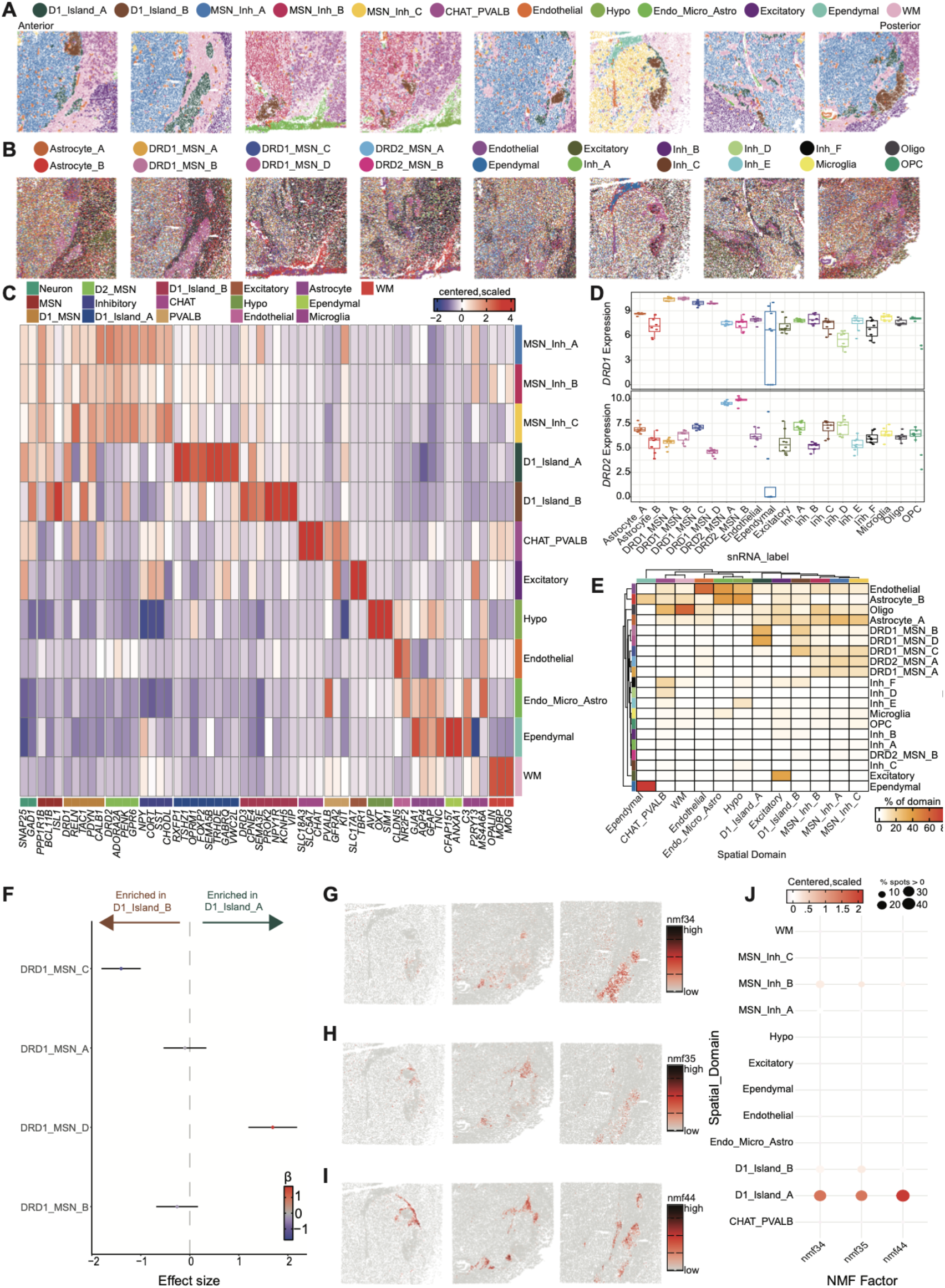
Topography and cell type composition of D1-islands with VisiumHD. **A**. Spatial clustering with *BANKSY* identified 12 spatial domains, including two populations of *DRD1* expressing D1-islands. **B**. Tissue sections with cells colored by predicted cell type from published NAc snRNA-seq data^2^ using label transfer. **C**. Expression profiles of 12 VisiumHD spatial domains. **E**. Heatmap depicting the percentage of predicted NAc cell types identified within each VisiumHD spatial domain. **F**. Compositional analysis identified DRD1_MSN_D as preferentially enriched in D1_Island_A and DRD1_MSN_C as preferentially enriched in D1_Island_B. Tissue arrays colored by loading for NMF patterns **G**. NMF34, enriched in DRD1_MSN_B, **H**. NMF35, enriched in DRD1_MSN_B, and **I**. NMF44, enriched in DRD1_MSN_D. **J**. Dotplot summarizing enrichment of island specific NMF patterns in each spatial domain.

To complement *BANSKY* clustering and enhance cell type resolution, we predicted cell type identity of each segmented cell using label transfer with our previously generated NAc snRNA-seq dataset^2^ (**Fig. 3B, Fig. S10**). This approach identified single MSN subtypes as shown by increased expression of *DRD1* in DRD1_MSN_A-D (**Fig. 3D**) and *DRD2* in DRD2_MSN_A-B (**Fig. 3D**) and is useful in understanding heterogeneous cell mixtures within spatial domains. To better understand the cellular organization of D1-island spatial domains, we quantified transcriptionally distinct cell types within each spatial domain (**Fig. 3E**). As expected, the WM domain contained cells almost exclusively predicted to be oligodendrocytes and astrocytes, while the MSN_Inh spatial domains contained a mixture of DRD1_MSN_A, DRD2_MSN_A, and DRD1_MSN_C cell types (**Fig. 3E**). D1_Island_A was made up of DRD1_MSN_B and DRD1_MSN_D, two transcriptionally distinct cell types that were previously mapped to D1_Islands within the NAc^2^. D1_Island_B was similarly enriched for transcriptionally distinct D1 subtypes, but also showed enrichment of DRD1_MSN_C (**Fig. 3E**). We quantified differences in composition using *crumblr*^25^ and found that cells predicted as DRD1_MSN_A and DRD1_MSN_B were similarly enriched within both VisiumHD D1-island spatial domains. In contrast, cells predicted as DRD1_MSN_C were preferentially enriched in D1_Island_B, while cells predicated as DRD1_MSN_D were preferentially enriched in D1_Island_A (**Fig. 3F**). To further investigate differential cell type enrichment across the island subtypes, we leveraged non-negative matrix factorization (NMF)^26^ patterns that were previously computed on human NAc snRNA-seq data^2^ to identify patterns of gene expression across the tissue that were independent of cell type and spatial clustering. We previously identified three NMF patterns associated with D1-islands that were highly correlated with transcriptional signatures of DRD1_MSN_B and DRD1_MSN_D cell types. In particular, NMF34 and NMF35 were highly enriched for DRD1_MSN_B, while NMF44 was highly enriched for DRD1_MSN_D, suggesting heterogeneity among D1-islands. To directly demonstrate enrichment of specific cell types in D1-islands, we projected these three NMF patterns into the VisiumHD dataset (**Fig. 3G-I, Fig. S11**). Further supporting preferential enrichment of DRD1_MSN_B and DRD1_MSN_D in the D1_Island_A spatial domain, NMF34, NMF35, and NMF44 were all enriched within the D1_Island_A spatial domain, with minimal signal in the D1_Island_B spatial domain (**Fig. 3J**). The relative depletion of NMF factors specific to canonical NAc D1-islands within the D1_Island_B spatial domain supported our conclusion that this domain represented human ICj.

### 2.4 Transcriptional signatures and molecular diversity of D1-island subtypes

To orthogonally validate D1_Island_A and D1_Island_B subtypes, we compared gene expression and spatial localization across the Xenium and VisiumHD data. Across platforms, we identified the same D1-island subtypes (**Fig. 4A**), which localized to the same topographic locations (**Fig. 1B, Fig. 4B**). Comparison of D1-island subtypes showed that *RXFP1* and *SEMA5B* are highly specific to D1_Island_A, while *VIP* and *PROK2* are highly specific to D1_Island_B (**Fig. 4C**). Using RNAscope single molecule fluorescence *in situ* hybridization (smFISH) to investigate the specificity of these marker genes, we confirmed the expression patterns identified within the VisiumHD dataset by showing minimal overlap between *PROK2*/*SEMA5B* and *VIP*/*RXFP1* (**Fig. 4D-I, Fig. S12**). These experiments provide tertiary evidence of *RXFP1*/*SEMA5B* as selective marker genes for D1_Island_A and *PROK2*/*VIP* as selective marker genes for D1_Island_B, which represent the ICj. The expression of *VIP* within ICj is in agreement with a recent publication that described a VIP expressing population of ICj-like cells.

**Figure 4.**
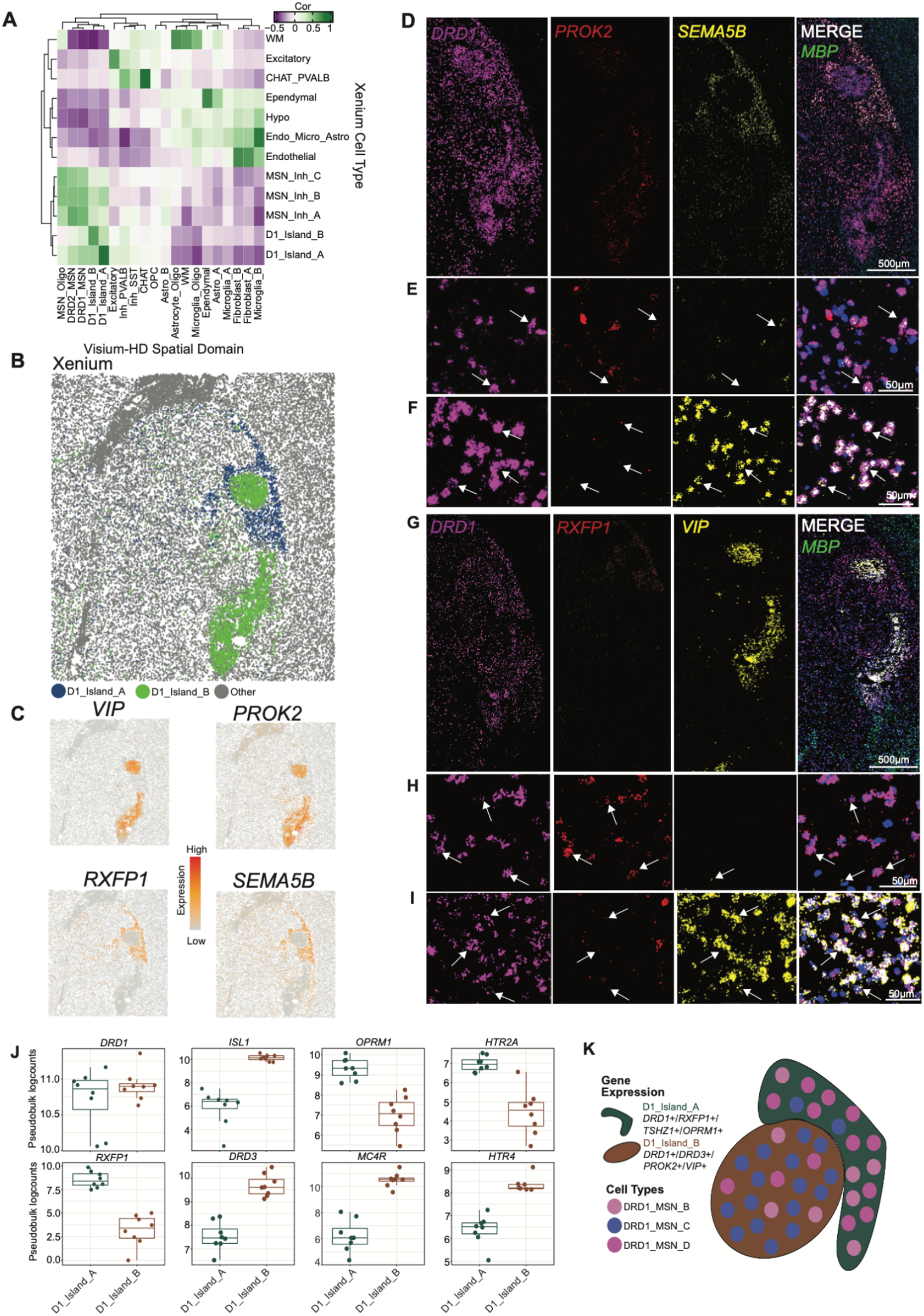
Identification of marker genes for transcriptionally distinct D1-islands in the human NAc. **A**. Heatmap depicting spatial registration results between Xenium cell types and VisiumHD spatial domains. **B**. Spatial plot zoomed in on D1-islands in a single Xenium array (Br6660, slice 8). **C**. Spatial feature plots depicting marker gene expression for D1_Island_A (*RXFP1*/*SEMA5B*) and D1_Island_B (*VIP*/*PROK2*). **D**. smFISH RNAscope of *DRD1, PROK2*, and *SEMA5B* within a single tissue section (Br6660, adjacent to slice 8). **E**. 40X images of *DRD1*+/*PROK2*+/*SEMA5B*-cells, and **F**. DRD1+PROK2-SEMA5B+ cells. **G**.smFISH RNAscope of DRD1, RXFP1, and VIP within a single tissue section. **H**. 40X images of DRD1+RXFP1+VIP-cells, and **I**. DRD1+RXFP1-VIP+ cells. **J**. Differences in expression of genes encoding peptides and receptors across transcriptionally distinct D1-islands. **K**. Diagram summarizing cell type composition and gene expression differences across D1-islands.

The identification of selective marker genes for D1-island subtypes is a first step towards molecular identification of these distinct cell types, which can facilitate future approaches for therapeutic targeting. To gain additional insight about potential biological functions of these cell types in the human brain, we examined expression of genes encoding neurotransmitter receptors and neuromodulatory peptides (**Fig. 4J**). D1_Island_A highly expressed *OPRM1* and *HTR2A*, the gene encoding the 5-HT2A receptor (**Fig. 4J**). Co-expression of *OPRM1* and*HTR2A* suggests that these cells may be responsive to specific drug classes, such as lysergic acid diethylamide (LSD), which acts on the 5-HT2A receptor, and opioids such as heroin,which act on the µ-opioid receptor. This is supported by findings in the rodent brain showing that D1-islands are modulated by opioids and regulate behavioral adaptations to these drugs^1,12^. D1_Island_B exhibited high expression of another serotonin receptor, *HTR4*, as well as *MC4R*, the gene encoding the melanocortin 4 receptor (**Fig. 4J**). *MC4R* is critically involved in feeding-related behaviors^27,28^ and the selective expression of this gene within D1_Island_B suggests that these cells may be involved in hedonic feeding. Together these experiments defined two transcriptionally distinct D1-island spatial domains distinguished by expression of *TSHZ1*/*RXFP1*/*OPRM1* and *DRD3*/*PROK2*/*VIP*, which contain preferential enrichment for DRD1_MSN_B and DRD1_MSN_D cell types in D1_Island_A and DRD1_MSN_C cell type in D1_Island_C (**Fig. 4K**).

### 2.5 D1-island subtypes are highly conserved across species

Molecular signatures of transcriptionally distinct MSN subtypes are highly conserved across rodents, NHPs, and humans^2,3,11^. Using MetaNeighbor^29^, a computational strategy that employs neighbor voting for identifying replicable cell types, we compared MSN subtypes across species and technologies with a focus on D1-islands (**Fig. 5A**). Hierarchical clustering revealed 3 primary categories consisting of D1-MSNs, D2-MSNs, and D1-islands (**Fig. 5A**). We found that NHP D2-Matrix neurons were transcriptionally similar to the DRD2_MSN_A cell type identified in human NAc snRNA-seq, and NHP D2-Striosome neurons were related to the human DRD2_MSN_B cell type (**Fig. 5A**). VisiumHD spatial domain DRD2_MSN was related to all D2-MSN cell types identified across species, demonstrating that this VisiumHD population likely contains several populations of transcriptionally distinct D2-MSNs. Within the rat and NHP striatum, heterogeneity within D1-MSNs was similar to that of D2-MSNs with different populations of D1-MSNs associated with NAc shell, striosome, or matrix (**Fig. 5A**). D1-islands make up the third primary category and split into two separate subcategories. The first category represented canonical NAc D1-islands marked by *DRD1*/*OPRM1*/*RXFP1*, and included NHP D1-NUDAP cells, rat Chst9-MSNs, rat Sema5a-MSNs, and D1_Island_A from human Xenium and VisiumHD. The second category of D1-islands represented ICj marked by *DRD3* and included NHP D1-ICj cells, rat ICj-Drd3 cells, and D1_Island_B from human Xenium and VisiumHD data, providing further evidence that human D1_Island_B represents ICj. The identification of ICj within the dorsal NAc was unexpected as these cells have primarily been characterized within the OT^19–21,23^.

**Figure 5.**
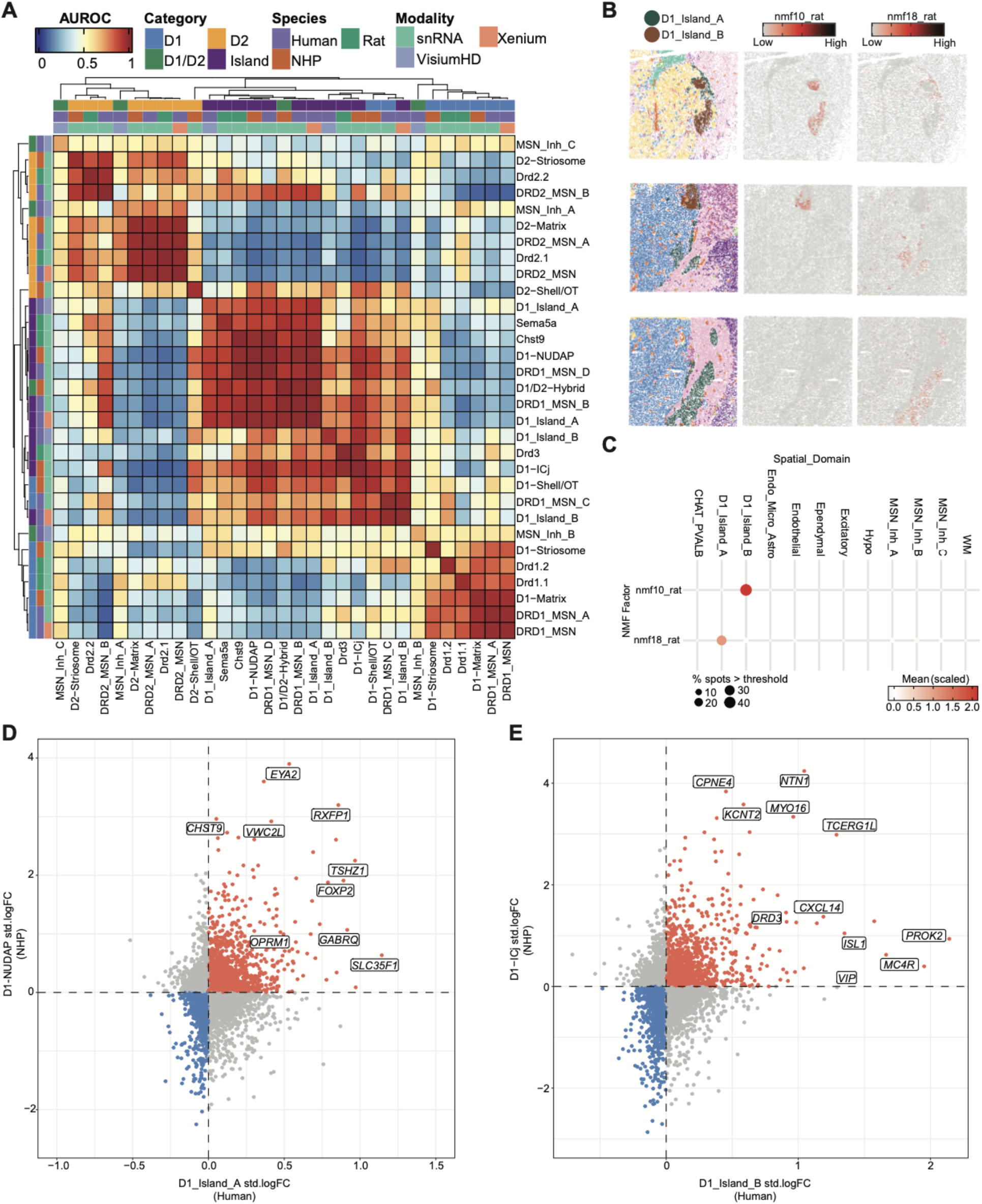
Molecularly distinct D1-island subtypes are conserved across species. **A**. Heatmap depicting MetaNeighbor AUROC scores across modalities and species. **B**. Non-negative matrix factorization (NMF) patterns specific to D1-islands and Islands of Calleja (ICj) identified in rat snRNA-seq data1 projected into human VisiumHD data. **C**. Summary of enrichment of NMF patterns across VisiumHD spatial domains. **D**. Correlation of standardized log_2_ fold-change for marker genes across ICj in human Visium HD data (D1_Island_B) and NHP ICj snRNA-seq data. **E**. Correlation of standardized log_2_ fold-change for marker genes across NAc D1-islands in human VisiumHD data (D1_Island_A) and NHP snRNA-seq data (D1-NUDAP).

We next aimed to understand cross-species conservation of human and rodent ICj. To do this, we used a rat snRNA-seq dataset^1^ containing a cluster of ICj cells marked by *Drd3*. We performed NMF on rat snRNA-seq data and identified two factors associated with both NAc D1-islands and ICj (NMF10 and NMF18) (**Fig. S13A-B**). To assess the spatial enrichment of NMF patterns, we projected these NMF patterns onto mouse Xenium data^1^. NMF10, which was associated with rat ICj cells, was highly enriched in the spatial location where ICj are found in the mouse brain (**Fig. S13C**). We then projected the rat NMF patterns into human VisiumHD data and found that ICj pattern NMF10 is enriched in D1_Island_B (**Fig. 5B,C**). NMF18, associated with a subpopulation of canonical D1-islands in the rat, was enriched in D1_Island_A (**Fig. 5B**,**C**). The enrichment of NMF patterns associated with rat D1-islands within human D1-islands further demonstrates the high degree of conservation of these cells across species.

Finally, we sought to understand conserved marker genes by correlating the standardized log fold-changes of 1-to-1 orthologs across humans D1-islands and their corresponding cell type in NHP (**Table S2, Table S3**).

Correlation of gene enrichment between human D1_Island_A and NHP D1-NUDAP identifies *FOXP2, RXFP1, VWC2L, TSHZ1, GABRQ*, and *OPRM1* as conserved marker genes for non-ICj canonical NAc-enriched D1-islands (**Fig. 5D**). *CHST9* was highly enriched within NHP D1-NUDAP cells, but only slightly enriched within the D1_Island_A spatial domain (**Fig. 5D**). Importantly, *CHST9* was differentially enriched between human snRNA-seq DRD1_MSN_B and DRD1_MSN_D, the two cell types that primarily reside in the D1_Island_A spatial domain (**Fig. 3E**). ICj are immature granule cells and the enrichment of *ISL1*, a developmentally important transcription factor, consistent with the fact that they contain immature granule cells. In addition to *DRD3* and *ISL1*, conserved markers of ICj include *CXCL14, NTN1, CPNE4, KCNT2, MYO16*, and *TCERG1L* (**Fig. 5E**). ICj extended into the OT, where *CXCL14* was expressed within OT protrusions ^30^. *CPNE4* is a specific marker of NHP ICj cells, but in human NAc, we find that *CPNE4* is more widely expressed across transcriptionally distinct D1-MSNs. This difference suggests that *CPNE4* may be a spatially variable gene that is enriched within human ICj cells, but is not a selective marker on the level of transcriptionally distinct D1-MSN cell types. Compared to D1_Island_A, *MC4R* was highly enriched within the D1_Island_B spatial domain, but only slightly enriched within the NHP D1-ICj population (**Fig. 5E**). Together, these results suggest a remarkable level of conservation between NHP and human island populations.

### 2.6 Dynamic co-localization of cell types along the AP axis in Xenium

Next, we used *CRAWDAD*^31^ to examine whether local cellular neighborhoods around D1-islands differed across the NAc AP axis by quantifying directional spatial relationships between reference and neighboring cell types within aligned Xenium sections. *CRAWDAD* is a reference-based tool that compares the local neighborhood of a reference cell type against a null background consisting of increasingly larger tile sizes with shuffled cell type labels. To evaluate whether local microenvironments differed across the NAc, we projected six Visium HD tissue footprints onto the aligned Xenium cells (**Fig. S14**) and used them to divide the tissue into lateral, dorsomedial, and ventromedial regions of interest (ROIs) (**Method 4.9, Fig. 6A, Fig. 6B**). We first summarized directional spatial relationships among all annotated cell types within the dorsomedial ROI of a representative Xenium slice (Br6660 slice 8) from an intermediate-to-posterior level. The resulting relationship matrix identified three groups of cell types that were consistently colocalized across the evaluated spatial scales (**Fig. 6C**). The first group comprised Fibroblast_A, Fibroblast_B, and Microglia_B and exhibited particularly strong positive spatial associations, consistent with a spatially restricted stromal and immune-associated compartment that is distributed across the tissue section. The second group contained D1_Island_A, DRD1_MSN, DRD2_MSN, MSN_Oligo, and Astro_B, and the third group contained D1_Island_B, WM, Astro_A, and Astrocyte_Oligo. Spatial visualization showed that the second and third groups occupied distinct tissue compartments, consistent with different local cellular environments around D1_Island_A and D1_Island_B within this section–ROI combination (**Fig. 6D**).

**Figure 6.**
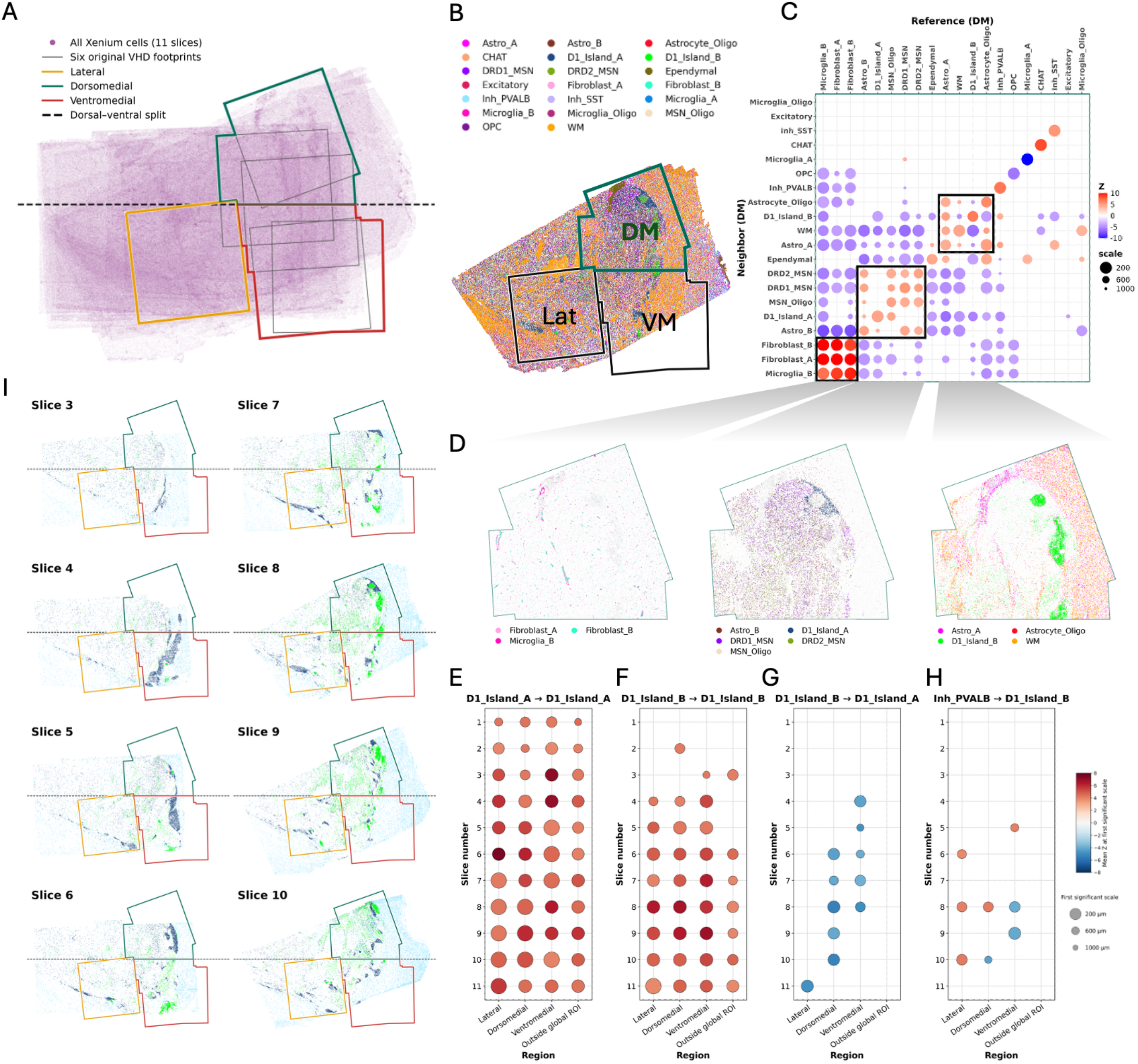
Spatial organization of cell types across anatomical regions and tissue depth in 11 aligned Xenium slices from Br6660. **A**. Six Visium HD tissue footprints projected onto the aligned Xenium cells and used to define the lateral (Lat), dorsomedial (DM), and ventromedial (VM) regions of interest (ROIs). **B**. A representative Xenium slice (Br6660 slice 8) illustrating the spatial distribution of annotated cell types, overlaid with three ROIs. **C**. Summary visualization of all cell type spatial relationships in the DM on slice 8. Point size represents the first significant scale (um), defined as the smallest evaluated shuffle scale at which the absolute mean Z-score reaches 3.84. Selected groups of consistently colocalized cell types are outlined by black boxes. **D**. Spatial visualization of the selected consistently colocalized cell types in the DM on slice 8. **E, F, G, H**. Summary visualization of four pairs of reference-neighbor cell type pairs in *CRAWDAD* analysis across slices and regions. Point size represents the first significant scale, defined as the smallest evaluated shuffle scale at which the absolute mean Z-score reaches 3.84. **I**. The distributions of D1_Island_A, D1_Island_B, and Inh_PVALB among Xenium tissue slices across anatomical depth and ROI.

We next compared the cell type relationship matrices across anatomical ROIs and tissue depths. Within the same tissue section, patterns of spatial enrichment and depletion differed among the lateral, dorsomedial, and ventromedial ROIs (**Fig. S15A**). Similarly, within the same ROI, anterior and posterior sections exhibited different spatial relationship profiles (**Fig. S15A**). These comparisons indicated that local cell type organization varied with both ML/DV ROIs and AP position. We next examined self-associations and directional relationships between the two D1-island populations across sections and ROIs. D1_Island_A showed self-enrichment across most evaluated AP levels and ROIs (**Fig. 6E**). In contrast, D1_Island_B self-enrichment emerged primarily at intermediate-to-posterior AP levels and was strongest within the dorsomedial and ventromedial ROIs (**Fig. 6F**). These positive self-associations supported the spatial aggregation of both populations into island-like domains rather than uniform cellular distributions (**Fig. 6I**). The AP distribution of D1_Island_B self-enrichment was consistent with its more restricted distribution in the smoothed 3D visualization (**Fig. 2D**). At intermediate-to-posterior AP levels, D1_Island_B reference cells showed consistent spatial depletion of neighboring D1_Island_A cells within the dorsomedial and ventromedial ROIs, with fewer D1_Island_A cells in their local neighborhoods than expected under the spatial null model (**Fig. 6G**). Although the populations occupied nearby anatomical territories at a broader spatial scale (**Fig. 6I**), this pattern was consistent with adjacent but locally segregated island domains rather than intermingled populations. No consistent corresponding pattern was observed when D1_Island_A was used as the reference population (**Fig. S15B**). This directional asymmetry may reflect differences in spatial extent: D1_Island_B occupied a more restricted domain, whereas D1_Island_A was more broadly distributed and only a subset of its cells lay near D1_Island_B. Finally, the directional relationship from a canonical GABAergic neuron subtype (Inh_PVALB) reference cells to neighboring D1_Island_B cells shifted between spatial enrichment and depletion across AP levels and ROIs (**Fig. 6H**). This suggested that the local positioning of these interneurons relative to D1_Island_B cells was anatomically context dependent rather than characterized by a uniform spatial association.

### 2.7 D1-islands are enriched for genes associated with risk for psychiatric disease and express targets of opioids, psychedelics, and antipsychotics

To better understand the potential functional relevance of the spatiomolecular structures and cell types we identified, we assessed enrichment for genes associated with genetic risk for brain disorders. Using the single cell disease risk score^32^ (scDRS), we tested the association between genetic risk for brain disorders and cell type-specific or domain-specific molecular signatures, analyzing VisiumHD spatial domains and cell type labels separately. We focused on GWAS for psychiatric disease^33–41^, substance use disorders and related traits^42–44^, cognition associated traits^39,45^, neurological disorders^46,47^, and included type 2 diabetes^48^ as a control. scDRS identified that D1_Island_B, which corresponds to human ICj, is enriched for genes associated with genetic risk for anorexia nervosa and (AN) and smoking cessation (SMK_CES, **Fig. 7A**). While not meeting statistical significance after adjusting for multiple testing, we also noted enrichment for cigarettes per day and smoking initiation (SMK_INIT) in D1_Island_B (**Fig. 7A**). Shared genetic risk for smoking-related traits and anorexia is interesting because increased nicotine use and dependence has been noted in patients with eating disorders^49,50^.

**Figure 7.**
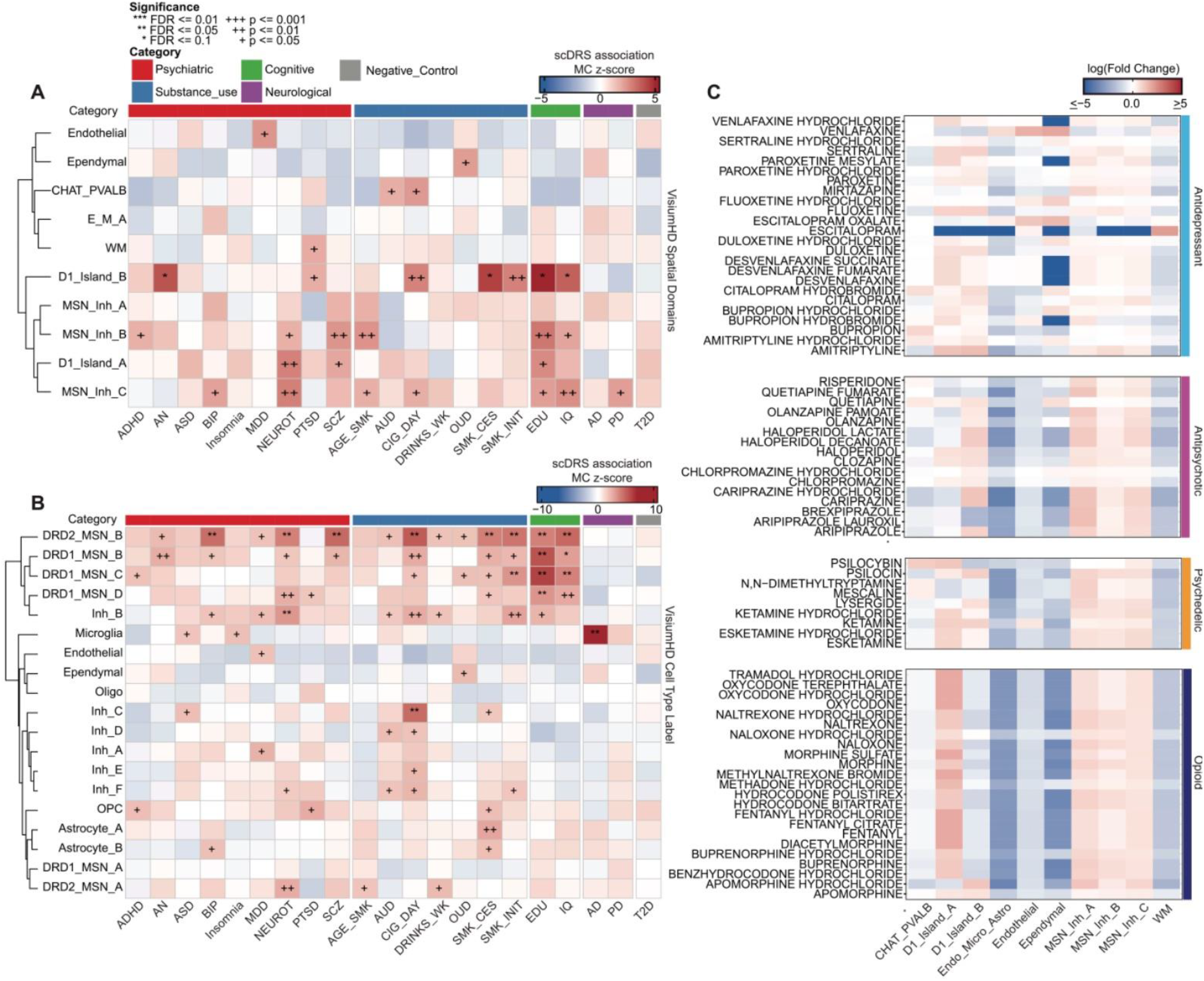
Genetic risk and drug targeting across NAc spatial domains and cell types. Heatmaps depicting scDRS association z-scores for brain diseases across **A**. spatial domains, and **B**. cell type labels. **C**. Heatmap depicting enrichment of genes encoding targets for antidepressants, antipsychotics, psychedelics, and opioids.

Because spatial domains may contain heterogeneous cell populations, we next asked which cell types within each spatial domain were driving enrichment of genetic risk for brain disorders. While genetic risk for psychiatric disease was enriched in some spatial domains, these structures may contain heterogeneous cell populations that could dilute genetic signals in some cases. Thus, we next investigated enrichment within discrete NAc cell types. Various D1- and D2-MSN subtypes (DRD1_MSN_B, DRD1_MSN_C, DRD1_MSN_C, and DRD2_MSN_B) are enriched for genes associated with genetic risk for both IQ and education (**Fig. 7B**). DRD1_MSN_C, a D1-MSN subtype primarily found in the D1_Island_B spatial domain (**Fig. 3F**), is enriched for genetic signals associated with smoking cessation (**Fig. 7B**), suggesting that enrichment identified in D1_Island_B may be driven by a single cell type. DRD2_MSN_B, but not DRD2_MSN_A, was also enriched for genetic signals associated with schizophrenia (**Fig. 7B**). *DRD2* is an established schizophrenia gene with the short isoform representing a potential causal isoform^51^. Additionally, DRD2 is the primary target of FDA-approved antipsychotics. Since *DRD2* is expressed across several NAc MSN subtypes, the enrichment of schizophrenia risk specifically in DRD2_MSN_B suggests that this subtype may represent a novel cellular target for the treatment of the disorder. However, we do not have the resolution to disentangle *DRD2* isoform expression and specifically link *DRD2* expression in this cell type to schizophrenia risk. Given that the ***µ***-opioid receptor and molecularly distinct NAc cell types are necessary for in opioid-related behavior in rodents ^1^, it was surprising that genetic risk for opioid use disorder (OUD) was not enriched in any specific spatial domain or cell type. The lack of genetic signal may reflect limitations of the underlying GWAS, as studies of substance use disorders are often underpowered due to societal stigma and complicated by comorbidity with a variety of other neuropsychiatric disorders. It is important to note that the enrichment of risk genes within a cell type or spatial domain does not, by itself, establish its role in cellular function or disease. It is important for follow up studies to provide experimental evidence for the role of risk genes in cellular function, as well as how disease-associated variants may affect normal function.

We next sought to understand which spatial domains may be engaged by different drug classes. Specifically, drug2cell^52^ was used to identify spatial domains containing expression of known targets of antidepressants, antipsychotics, psychedelics, and opioids. Drug2cell identified antidepressant targets as broadly enriched across both D1-island domains (**Fig. 7C**). D1_Island_A, which contains molecularly distinct populations of *OPRM1*-expressing D1-MSNs, were highly enriched for targets associated with different classes of opioids, including fentanyl, oxycodone, and morphine (**Fig. 7C**). This result is supported by recent evidence demonstrating that cell types within rat D1-islands are directly engaged by opioid receptor agonists ^1^. D1_Island_A was also enriched for targets associated with psychedelics, including psilocybin (**Fig. 7C**). This is highlighted by high expression of *HTR2A* within the D1_Island_A spatial domain (**Fig. 4J)**, which is directly engaged by psilocybin. The enrichment of targets for opioids and psychedelics suggests that cell types within D1_Island_A may represent a novel cellular target for treating substance use disorders, and other disorders where psychedelics are being considered as potential therapies. Although additional work to identify targetable molecules within this cell type are needed to mimic therapeutic effects of psychedelic drugs. MSN_Inh_A, MSN_Inh_B, and MSN_Inh_C,, are enriched for targets of antipsychotics (**Fig. 7C**). These spatial domains contain D2-MSNs, which are cellular targets for treating psychosis in schizophrenia and bipolar disorder. D1_Island_B, corresponding to human ICj, exhibits high *DRD3* expression (**Fig. 4J)**, and enriched for targets of cariprazine, aripiprazole, and brexpiprazole (**Fig. 7C)**, antipsychotics that engage the *DRD3* receptor and position human ICj as a cellular target for treatment of schizophrenia. Altogether, the drug2cell results highlight the potential ability to target unique molecular structures within the NAc for the treatment of brain disease.

## 3 Discussion

Spatial transcriptomics studies have substantially expanded our understanding of the molecular and cellular architecture of the human brain. However, most studies have profiled a single or limited number of 2D tissue sections, and therefore cannot capture variation in cell type abundance, spatial topography and local cellular organization across the 3D structure of a given brain region. This limitation is particularly relevant in the NAc, where afferent connections exhibit topographic organization across anatomical axes and molecularly distinct neuronal populations occupy characteristic spatial locations By systematically profiling the entire AP extent of the human NAc, we generated a 3D spatiomolecular reconstruction, and found that although the topographic location of major NAc neuronal populations was largely maintained across the AP axis, their relative abundance and local cellular environments varied with anatomical position. These differences were particularly pronounced for D1-islands, which we resolved into two molecularly distinct populations with different 3D distributions and local spatial relationships. Together, these findings demonstrate that AP position is an important dimension of molecular neuroanatomical organization that is not routinely captured by sampling 2D tissue sections.

Our previous work suggested that the abundance of D1-islands varies across the AP axis, with greater representation at intermediate AP levels; however, because AP position was sampled inconsistently across different donors, anatomical position was confounded with inter-individual variability. By systematically sampling the AP axis within individual donors, the current study demonstrates that D1-island abundance varies with AP position, and further resolves this variation across two molecularly distinct island populations. D1_Island_A, marked by *RXFP1, OPRM1*, and *GABRQ*, is distributed throughout the AP axis, with greater representation at anterior-to-intermediate levels, whereas D1_Island_B, marked by *DRD3, VIP, PROK2*, and *ISL1*, is more restricted to intermediate-to-posterior levels. Across AP levels, both populations maintain characteristic spatial distributions and organize into discrete bundles along the NAc borders.

Most MSN populations within the human NAc exhibit overlapping spatial distributions organized along continuous gradients rather than forming sharply delineated anatomical domains^2^. D1-islands represent a notable exception, organizing into discrete, densely packed bundles of cells along the medial and ventral-lateral border of the NAc. Our analysis of local cellular microenvironments further supports this distinct spatial organization, demonstrating that both D1-island domains consist of these local cell clusters. Despite occupying adjacent anatomical locations, D1_Island_A and D1_Island_B also exhibit local spatial segregation, with D1_Island_A cells depleted from the local microenvironment surrounding D1_Island_B at intermediate-to-posterior AP levels. Studies across the rodent^1,6,11^, NHP^8^, and human^2^ have identified a similar phenomenon with D1-islands forming a discrete spatial domain of tight bundles along the medial and ventral borders of the NAc. Moreover, relationships between D1-islands and other neuronal populations were not uniform across the NAc. For example, the spatial relationship between paravalbumin-exressing GABAergic neurons and D1_Island_B shifted between enrichment and depletion depending on AP position and dorsomedial or ventromedial location. Together, these findings indicate that although D1-islands maintain a distinct spatial organization, the local cellular environments in which they are embedded varies across the 3D anatomy of the NAc. These findings are broadly consistent with a recent spatial analysis of the human striatum that identified molecular gradients across dorsal and ventral striatal regions ^7^. However, by densely sampling within the NAc across the AP axis, our study resolves finer-scale variation in the abundance and local organization of D1-island populations that are not captured by mesoscale analyses across the broader striatum.

Several recent studies have demonstrated substantial conservation of molecular signatures across NAc MSN populations in rodents, NHPs and humans^2,3,5,12,53^, with D1-islands exhibiting conservation of both molecular identity and topographical organization. Here, we resolved two molecularly distinct island populations that map to conserved populations across species. D1_Island_A corresponds to previously described NAc D1-islands, including NHP D1-NUDAP^8^ and rat Sema5a/Chst9-MSNs^1,11^, and shares conserved expression of *RXFP1* and *OPRM1*. In contrast, multiple lines of cross-species evidence identify D1_Island_B as human ICj, including its molecular similarity to both NHP D1-ICj and rodent ICj populations as well as conserved expression of *DRD3* and *ISL1*. While ICj have primarily been characterized within the olfactory tubercle in the rodent and NHP brain, in the human brain we found these cells to extend along the entire medial border of the NAc and into the dorsal aspects of the NAc proper. One possibility is that some of these more dorsally positioned cells, are related to the insula magna, a well-characterized ICj population located near the the NAc-septal border ^8^ in rodents. Regardless, the distribution of D1_Island_B within the human NAc differs from the classically described organization of rodent ICj, suggesting that the spatial organization of this conserved population may differ across species. Molecular differences were also evident, including prominent *VIP* expression within human ICj that was not similarly conserved across species. *VIP* expression within human ICj has also been validated in another human atlas of NAc^53^. Thus, although both island populations exhibit substantial conservation of molecular identity across rodents, NHP, and humans, their molecular features and anatomical organization also reveal species-specific specialization.

A major limitation of post-mortem human molecular studies is the inability to directly test the function of identified cell populations. However, molecular conservation across species provides an opportunity to generate hypotheses about the functions of homologous populations in the human brain. D1_Island_A is marked by *TSHZ1* and *OPRM1*, and corresponds to a conserved rodent D1-island population that has been functionally studied by targeting Chst9- and Tszh1-expressing MSNs^11,12^. In the rat, Chst9-MSNs are directly inhibited by μ-opioid receptor activation, and selective deletion of *Oprm1* in these cells prevents fentanyl-conditioned place preference, demonstrating a requirement for μ-opioid receptor signaling in this population for opioid reward learning. *Tshz1*-expressing MSNs have also been implicated in opioid withdrawal in the mouse NAc, where their activation locally inhibits DA release^12^. Deletion of *Oprm1* from *Tshz1*-expressing neurons blunts both the reduction in NAc DA release and the aversive response associated with opioid withdrawal. Together, these findings suggest that the conserved D1-island population represented by D1_Island_A may be particularly important for opioid-dependent modulation of NAc circuitry in the human brain. This hypothesis is further supported by our previous finding that gene expression patterns associated with volitional morphine intake are enriched within human D1-islands^2^. In contrast, considerably less is known about the function of ICj, although recent studies have implicated these cells in orofacial grooming^19^ and depressive-like phenotypes^20^. Together with their distinct receptor and neuropeptide expression profiles, the molecular conservation of ICj across species provides a framework for testing whether functions identified in model systems extend to the corresponding human population. In summary, the molecular correspondences identified here provide experimentally tractable cross-species targets for investigating the functions of these spatially organized NAc populations in the human brain.

A complementary method to infer function and understand potential behaviors regulated by human cell types is to investigate enrichment of genetic risk for disease. Using scDRS, we show that human ICj (D1_Island_B) is enriched for genetic signals associated with anorexia nervosa and traits associated with smoking. While some research has been dedicated to the molecular profiling of ICj across lower-order mammals, little research has focused on the role of these cells in complex behavior. Recent work suggests they are important for orofacial grooming and depressive behaviors^19,20^. The enrichment for genetic signals associated with anorexia nervosa and smoking motivates generation of hypotheses to test whether this structure is directly involved in feeding and addiction-related behaviors. In addition to harboring genetic risk for several diverse brain disorders, D1-islands are also enriched for molecular targets of antipsychotics, psychedelics, and opioids. The enrichment of targets for opioids within D1-islands is particularly interesting because *Oprm1* expression in D1-islands is required for fentanyl-induced reward learning in rodent models of behavior^1^. Further, the co-enrichment of targets for opioids and psychedelics suggests that D1-islands may represent a cellular target for psychedelic-assisted therapies for opioid use disorders (OUDs), which are actively being investigated as an OUD therapy^54^.

The experiments here were designed as a deep anatomical profiling study, prioritizing dense within-donor sampling across the entire extent of the AP axis in the human NAc. This design enabled reconstruction of molecular and cellular organization across three dimensions, but was limited to 2 neurotypical donors and therefore cannot establish the extent of inter-individual variation in these patterns or their relationships with sex, age, or disease. Larger studies will be necessary to determine the generalizability of the spatial features identified here. Comprehensive AP sampling also critically depends on the orientation and anatomical integrity of the postmortem human tissue blocks, which are prone to freezing induced alterations in morphology. Although Br6660 exhibited well-preserved anatomy across the AP axis, Br6436 was affected by an angled coronal slab, which complicated anatomical registration at posterior levels, and consequently the most detailed 3D analyses focused on Br6660. Finally, Xenium profiling was performed using a targeted 366-gene panel designed to distinguish major NAc neuronal populations, particularly focusing on D1-island markers. While this design enabled high-resolution characterization of D1-island heterogeneity, it did not capture the full transcriptional diversity of D1- and D2-MSN populations previously identified by transcriptome-wide approaches. VisiumHD profiling provided complementary near-transcriptome-wide characterization of selected ROIs, but at substantially fewer anatomical positions. Future studies combining dense 3D sampling with broader transcriptomic coverage and larger donor cohorts will be necessary to determine how the spatiomolecular organization identified here varies across individuals and disease states.

In conclusion, systematic profiling across the extent of the human NAc revealed molecularly distinct cell populations and spatial features that would be missed, or substantially undersampled by analysis of a single 2D section. By resolving changes in cell type proportion and local cellular organization across three dimensions, this study highlights the importance of anatomical position for interpreting the molecular architecture of the human NAc. In particular, we identified two molecularly distinct D1-island populations with different 3D distributions, one of which we unexpectedly revealed as ICj, providing evidence that in the human brain this population extends into the NAc beyond its classically described location in the OT. The molecular conservation of these populations across species provides a framework for linking cell identity and spatial topography in the human brain to circuit and functional studies in experimentally tractable model systems. Future studies integrating 3D molecular organization with projection-defined connectivity will be important for determining how these spatially-organized neuronal populations are incorporated into NAc circuits that support reward-related behavior and are altered in neuropsychiatric and substance use disorders.

## 4 Methods

### 4.1 Postmortem human tissue samples

Postmortem human brain tissue from neurotypical adult donors of European ancestry (*N* = 2) were obtained at the time of autopsy following informed consent from legal next-of-kin through the Maryland Department of Health IRB protocol #12–24. Using a standardized strategy, all donors were subjected to clinical characterization and diagnosis. Macroscopic and microscopic neuropathological examinations were performed, and subjects with evidence of significant neuropathology were excluded. Additional details regarding tissue acquisition, processing, dissection, clinical characterization, diagnoses, neuropathological examination, RNA extraction and quality control (QC) measures have been previously published^114^. The two control donors had no lifetime history of psychiatric or substance use disorders according to DSM5 and were negative for acute alcohol and drug intoxication at the time of death. The male donor (Br6436) was 58.4 years old at the time of death and was of Caucasian descent, with postmortem interval (PMI) of 37.5 hours and best RNA integrity number (RIN) of 7.7 measured in the prefrontal cortex (PFC). The female donor (Br6660) was 50.8 years old at the time of death and was of Caucasian descent, with PMI of 21 hours and best RIN of 7.5 measured in PFC. Tissue blocks (approximately 45 × 35 × 25 mm) containing the entire NAc in a single coronal fresh-frozen brain slab were selected for this study and were dissected using a hand-held dental drill. Tissue blocks were stored in sealed cryogenic bags at −80°C until cryosectioning.

### 4.2. Tissue processing and anatomical validation

Tissue slabs containing the entire NAc in one fresh-frozen slab were selected based on anatomical landmarks of the anterior and posterior faces of the slabs (**Fig. S1**,**Fig. S2**). The following landmarks were used to guide the brain block dissections: corpus callosum as the dorsal landmark, brain midline as the medial landmark, the orbitofrontal cortex as the ventral landmark, and the claustrum with surrounding white matter as the lateral landmark. Fresh frozen tissue blocks were acclimated to −14^°^C for 30 minutes inside the cryostat (Leica CM3050s), mounted on an extra large round chuck with Optimal Temperature Compound (TissueTek Sakura, Cat #4583), and the tissue was trimmed until the appearance of a clear ventral striatum from the anterior face of the block. Several 10 µm tissue sections were then collected on pre-chilled glass microscope slides (VWR SuperFrost Microscope Slides, Cat #48311703) and banked at −80^°^C for later use. The tissue block was then scored with a razor blade to isolate the ventral striatum, a 10 µm section was mounted on a pre-chilled Xenium Spatial Gene expression slide (PN-1000460, 10x Genomics, Pleasanton, CA), and banked at −80^°^C for future Xenium processing. Subsequently ~400 µm of ventral striatal tissue was collected in a LoBind microcentrifuge tube (WVR, 80077-234) and banked at −80^°^C for future use. This cryosectioning procedure was repeated 10 additional times, resulting in 11 total Xenium slides taken ~500 µm apart per brain donor. All Xenium slides were processed in pairs within one month, to follow recommended storage procedures by the 10X Genomics (Pleasanton, CA). Following Xenium analysis, banked tissue sections were selected for Visium HD assay, specifically targeting regions containing D1-islands (Br6660: AP levels 4, 6, 7, 8, and 9; Br6436: AP levels 1 and 9). Following Visium HD analysis, additional banked tissue sections were selected for RNAScope validation experiments.

### 4.3 Xenium data generation

#### Xenium slide preparation

Xenium tissue slides were processed using the 10x Genomics (Pleasanton, CA) sample preparation and gene expression v1 workflows (CG000581 and CG000582, respectively). Procedures related to cryosectioning and slide storage were followed according to the demonstrated protocol CG000579. Briefly, banked tissue blocks were removed from −80°C storage and allowed to acclimate to −16°C in a CM3050 S Cryostat (Leica Biosystems). Simultaneously, 2 Xenium slides (PN-3000941) were removed from their mylar packaging at −20°C and were acclimated in the cryostat. The tissue was scored with a razor to isolate the NAc and one 10 µm tissue slice was mounted per Xenium slide. All sections were mounted within the marked sample area (10.45mm x 22.45mm) and slides were stored in a slide mailer at −80°C for 4-6 weeks before downstream fixation and permeabilization steps (CG000581).

Slides were processed in batches of two in anterior to posterior order within each donor. Slides were removed from −80°C and incubated on a Xenium Thermocycler Adaptor (PN-3000954) at 37°C for 1 minute and then fixed for 30 minutes in 40 mL of 3.7% formaldehyde (ThermoFisher Scientific, BP24731), washed in 40mL 1xPBS (ThermoFisher Scientific, AM9624) and permeabilized in 1% SDS solution (Millipore Sigma 71736) diluted in 0.2 µm filtered nuclease-free water (IDT 11-05-01-04) for 2 minutes. Following removal of the SDS solution, the slides were washed 3x (40 mL/wash) in 1xPBS and then immersed in 40 mL chilled 70% methanol (Millipore Sigma 34860) diluted in nuclease-free water. After 60 minutes, slides were removed and rinsed 2x in 1xPBS (40 mL/wash) and then placed in the Xenium Cassettes (PN-1000566). To complete the subsequent probe hybridization, ligation, and amplification steps, we followed procedures outlined in CG000582. To accommodate the 20 reactions needed for this study, we purchased and resuspended a 16rxn and a 4rxn Xenium Add-On Custom 51-100 gene panel (PN-1000561 or PN-1000651, respectively) in 700 µL or 140 µL TE Buffer respectively (ThermoFisher Scientific BP24731). The custom probes were then resuspended in Xenium Probe Hybridization Buffer (PN-1000390) containing the Xenium Human Brain Gene Expression Panel (PN-1000599) and applied to the Xenium slides. The probe hybridization mix was allowed to hybridize to accessible RNA during a 16-24hr overnight incubation at 50°C. The next day, hybridization reactions were washed 2x in 0.05% 1xPBS-Tween-20 (Tween-20: Thermo Fisher Scientific 28320) (500 µL/wash) and then incubated in 500 µL Xenium Post Hybridization Wash Buffer at 37°C for 30 minutes. After 3 washes of 0.05% 1xPBS-Tween-20 (500 µL/wash), probes were ligated at 37°C for 2 hours in a reaction consisting of 87.5% Xenium Ligation Buffer (PN-2000391), 2.5% Xenium Ligation Enzyme A (PN-2000397) and 10% Xenium Ligation Enzyme B (PN-2000398). After 3 washes of 0.05% 1xPBS-Tween-20 (500 µL/wash), rolling circle products were amplified in a reaction consisting of 90% Xenium Amplification Mix (PN-2000392) and 10% Xenium Amplification Enzyme (PN-2000399) for 2 hours at 30°C. After amplification was completed, slides were washed 3x in TE buffer (500 µL/wash) and quenched for autofluorescence. During the quenching process, 500 µL of 1% Reducing Agent B (PN-2000087) diluted in 1xPBS was applied to the slides for 10 minutes at room temperature. Following 3 rinses ethanol (Millipore Sigma e7023) rinses (1 mL/rinse), slides were dehydrated for 5 minutes at 37°C and then rehydrated in 1 wash of 1xPBS followed by a 2-minute incubation of 0.05% 1xPBS-Tween-20. Nuclei were then stained in 500 µL Nuclei Staining Buffer (PN-2000762) and then washed 3x in 0.05% 1xPBS-Tween-20 (1 mL/wash). Slides were then stored overnight in the dark at 4°C and imaged on the Xenium Analyzer the following day.

#### Xenium Analyzer

Imaging of all slides took place on the Xenium Analyzer (PN-1000569) located at the Johns Hopkins Sidney Kimmel Comprehensive Center (SKCCC). Reagents and procedures needed to run the Xenium v1 workflow were followed according to the manufacturer’s instructions (CG000584). Software versions (v3.2.1.2) and onboard analysis versions (v3.2.0.7) were selected for Br6660 (slice number 1-11) and Br6436 (slice number 1-8). During the intervening scan between Br6436 slice number 8 and 9, a software update was implemented such that Br6436 slice number 9-11 were analyzed using the software and onboard analysis version 3.3.0.1. All runs passed a ‘readiness test’ confirming that the system was working optimally and that the instrument was ready for use. Reagents bottles necessary for imaging were prepared accordingly: Bottle 1 – Milli-Q ultrapure water (Thermo Scientific 50131948), Bottle 2 – 0.05% 1xPBS Tween-20, Bottle 3 – Milli-Q ultrapure water and Bottle 4 – 0.1% Tween-20, 50mM KCL (Teknova P0330), 50% DMSO (Millipore Sigma 41639) diluted in nuclease-free water (ThermoFisher Scientific AM9932). Xenium decoding consumables (PN-1000487) were loaded according to details provided by the manufacturer. Following successful loading of the instrument, panel ROI selection was performed and the run was initiated. Runs took on average of 48 hours to complete, after which consumables were unloaded and discarded. All images and supporting data were copied and transferred to the Lieber directory on the Johns Hopkins University Joint High Performance Computing Exchange (JHPCE).

### 4.4 Visium HD data generation

Tissue slides banked for Visium HD were processed in pairs using the Visium HD Spatial Gene Expression platform by 10x Genomics (Pleasanton, CA). Slides were removed from −80°C storage and tissues were fixed in pre-chilled methanol (Millipore Sigma, 322415), dehydrated in isopropanol (Millipore Sigma, I9516), and H&E-stained at room temperature, following the procedures adapted from CG000684 rev B. In brief, tissue was incubated in Gills II Hematoxylin (Leica, 3801520) for 1-minute, washed 3x in Milli-Q ultrapure water (800mL/wash) (Thermo Scientific, 50131948), then incubated in Bluing Buffer (Agilent, CS70230) for 1 minute and washed 3x in Milli-Q ultrapure water (800mL/wash). The tissue was then stained in Alcoholic Eosin (Leica, 3801615) and washed 3x in Milli-Q ultrapure water (800mL/wash). Slides were then cover-slipped in 85% glycerol (Acros Organics, 327255000) diluted in 0.2mm filtered nuclease-free water (IDT, 11-05-01-04) and 600U of Protector RNase inhibitor (Roche, RNAIHN-RO). Slides were imaged with a 40x 0.75 NA objective on a Leica Aperio CS2 digital pathology slide scanner (Leica Biosystems). Following coverslip removal, regions of interest (ROIs) for both tissue slides were selected via precise application of the Visium Cassette S3 gasket (CG000730 Rev A). ROIs were subsequently destained in 0.1N HCL (Fisher Chemical, SA54-1) for 15 minutes at 42°C and washed 3x in Tris-EDTA pH 8 (ThermoFisher Scientific, BP24731) followed by 1 wash of 1x PBS pH 7.4 (ThermoFisher Scientific, AM9624). Subsequent *In situ* hybridization, ligation, CytAssist-Enabled probe release, and library construction steps were followed according to the procedures outlined in CG000685 rev C. Briefly, a 15-minute permeabilization at room temperature in a 0.7% Tween-20 (Thermo Fisher Scientific, 28320) 1xPBS pH 7.4 solution, the Visium Human Transcriptome probe kit v2 (PN-1000466) was applied to each ROI and allowed to hybridize to accessible RNA at 50°C overnight. Hybridized tissues were then washed in FFPE Post Hyb Wash Buffer (PN-2000424) and ligated using a ligation enzyme (PN-2000425) diluted in 2x probe ligation buffer (PN-2000445) for 1 hour at 37°C. While tissues were washed in post ligation wash buffer (PN-2000419) and then allowed to acclimate to room temperature, a Visium HD slide (PN-1000670) was removed from −80°C storage warmed to room temperature for 30-60 minutes, and washed 3x in a total of freshly-prepared 60mL 0.1x SCC (Millipore Sigma S66391L). Additional washes were implemented, as needed, until both 6.5mm spatially-barcoded oligonucleotide capture arrays were visibly free of debris. The Visium HD slide was then placed in a Visium 2-port cassette S2 (PN-1000669) and RNase enzyme (PN-3000605) was allowed to equilibrate the arrays. During the Visium HD slide equilibration, the two tissue slides were stained in 10% Alcoholic Eosin, washed 3x in 1mL 1xPBS, and then mounted on the Visium CytAssist instrument (PN-1000441). After equilibration, the Visium HD slide was loaded in the CytAssist instrument, dried, and the probe release mix (PN-2000411, PN-3000605) was added to the center of each spacer well. The CytAssist run was then initiated. Immediately upon completion of the run, the Visium HD slide was washed 3x in Buffer EB (1mL/wash) (Qiagen, 19086) and placed in a Visium 2-port cassette. Extension enzyme (PN-2000389) and extension buffer (PN-2000409) were then applied to each capture array over two cycles of a 30min incubation at 53°C. Probes were eluted and then pre-amplified using AMP B (PN-2000567) and TS primer Mix B (PN-2000567) to generate ample material for downstream library construction. To determine the ideal number of additional cycles needed to amplify each library for the sample index PCR, pre-amplified samples were quantified using KAPA SYBR Fast qPCR Master Mix Universal (Roche KK4600) run on the CFX Opus 96 (Bio-Rad 12011319). Unique indices from the dual index plate TS Set A (PN-3000511) were assigned to each sample so that Visium HD libraries could be pooled and demultiplexed in downstream sequencing runs. CytAssist Spatial Gene Expression libraries were sequenced on the Illumina NovaSeq X Series 25B flow cell, targeting ~25000 reads per spot at Psomagen (Rockville, MD).

### 4.5 Xenium raw data processing and quality control

After imaging with the Xenium Analyzer, raw imaging data were processed using the Xenium Onboard Analysis (XOA) pipeline. XOA performs image processing, nucleus and cell segmentation, transcript decoding, duplicate removal, quality-score filtering, and assignment of decoded transcripts to segmented cells. The resulting cell-feature matrix was used as input for downstream quality control and preprocessing.

Quality control was performed on the Xenium cell-level count matrix stored as a *SpatialExperiment*^55^ object. We first removed cells with zero total counts across all features. We then calculated per-cell quality-control metrics using `addPerCellQCMetrics()` from *scuttle*^56^ v1.18.0, with subsets corresponding to gene-expression targets and technical control features. Technical control features included negative control codewords, negative control probes, and unassigned features. Cells with a high fraction of counts from negative-control or unassigned features were flagged as potential low-quality cells because these features reflect decoding uncertainty, nonspecific signal, or off-target background rather than targeted gene expression.

After computing control-feature metrics on the full feature set, we restricted downstream analysis to gene-expression panel features. Cells with no remaining gene-expression counts were removed. We then applied three cell-level quality-control filters. First, cells for which negative-control or unassigned features accounted for at least 25% of total counts were removed to reduce the influence of decoding errors and nonspecific signals. Second, cells with abnormally low total gene-expression counts or low numbers of detected gene-expression features were removed using sample-specific lower-tail outlier thresholds based on four median absolute deviations (MADs) on the log-transformed distributions. Third, cells with abnormal cell areas were removed using sample-specific two-sided four-MAD thresholds. This area-based filter was intended to exclude very small segmented objects, which may represent debris or fragmented cells, as well as very large segmented regions, which may represent merged cells or segmentation errors. We considered spatially aware quality-control procedures such as *SpotSweeper*^57^, which identifies local outliers and regional artifacts by comparing spatial transcriptomics quality metrics within local neighborhoods. However, we did not use *SpotSweeper* as the primary filtering strategy because it was designed for sequencing-based, spot-level spatial transcriptomics technologies such as Visium, whereas local outlier thresholds are less interpretable and can be unstable in heterogeneous cell-level Xenium neighborhoods.

After quality-control filtering, expression values were normalized using morphology-derived size factors rather than conventional library-size normalization. Unlike in scRNA-seq, total transcript counts in imaging-based spatial transcriptomics reflect targeted panel design, cell morphology, cell type, and local tissue biology rather than sequencing depth alone^58,59^. We therefore normalized by cell area to account for transcript density while preserving biologically meaningful variation associated with total expression. Specifically, nucleus-area size factors were computed by dividing each cell’s nucleus area by the median nucleus area across cells:

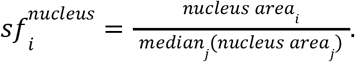

The raw count matrix was then normalized using `normalizeCounts()` from *scuttle*^56^ v1.18.0 with nucleus-area size factors and log_2_ transformation. Since cell and nucleus areas were highly correlated in this dataset, cell-area-derived size factors were also calculated during preprocessing as a sensitivity check. All downstream analyses used the nucleus-area-normalized expression matrix.

### 4.6 Xenium dimensionality reduction and clustering

#### Dimensionality Reduction

Dimensionality reduction was performed using *BANKSY*^60^ v1.4.0, a spatial clustering algorithm that generates a neighborhood augmented expression matrix that takes into account a cell’s gene expression and nearest neighbors. The augmented BANKSY matrix was calculated using k=30 nearest neighbors with `computeBanksy()`. Samples were processed individually for `computeBanksy()` and then merged with `cbind()` prior to dimensionality reduction. Following merging of the samples, principal component analysis (PCA) was performed with `runBanksyPCA()` with “Sample” as the grouping factor and seed=1000. The BANKSY workflow includes a hyperparameter, ***λ***, which determines how much spatial information to include. Our PCA was performed using a ***λ*** value of 0, which excludes nearest neighbor contributions and is based solely on gene expression^60^. Following PCA, uniform manifold approximation and projection (UMAP) was used to further reduce the dimensionality of the data for plotting purposes. UMAP was generated using `runBanksyUMAP()` with ***λ***=0 and `seed=1000`. The functions `computeBanksy()`, `runBanksyPCA()`, `runBanksyUMAP()`were run without use of the azimuthal gabor filter, which is a measure of gradients of gene expression within a cell’s neighborhood. Importantly, dimensionality reduction was performed within a single donor that exhibited well-preserved neuroanatomy. Batch correction was not applied so as to not erase any biological heterogeneity that may exist along different anatomical axes.

#### Clustering and cell type identification

Our first objective with the Xenium data was to ask whether transcriptionally distinct cell types exhibited unique topography across the dorsal-ventral, medial-lateral, and anterior-posterior axes. To do this, we performed clustering with functions provided by *BANKSY* v1.4.0. Clustering was performed within the PCA space with ***λ***=0, again excluding nearest neighbor information and instead clustering solely on gene expression information. Specifically, we performed louvain clustering with a resolution value of 0.5 using the `clusterBanksy()` function with `use_AGF=FALSE` and `seed=1000`, resulting in 22 distinct clusters or cell types. Importantly, clustering was performed within the tissue sections of a single donor that exhibited well-preserved neuroanatomy. This was the same donor in which dimensionality reduction was performed.

Our Xenium panel was designed to investigate neuronal heterogeneity, while allowing for the identification of non-neuronal cell types. Of the 22 identified clusters, 6 exhibited high expression of *OPALIN, MOG, MOBP, FGFR2*, and *PROX1*, genes that are highly enriched in white matter and oligodendrocytes^2^. Of these 6 cell types, 3 also exhibited high expression of genes enriched in MSNs, astrocytes, and microglia. These cell types likely include segmented cells that contain both an oligodendrocyte and another cell type. Based on this we called these cell types MSN_Oligo, Astrocyte_Oligo, and Microglia_Oligo. The other 3 cell types exhibited selective expression of oligodendrocyte genes and differed based on the levels of expression of these same genes. Additionally, these cells were found within white matter tracks that are located within the internal capsule and surrounding the striatum. Because our panel was not designed to investigate heterogeneity within white matter cells, we merged these 3 cell types into a single cluster called WM. We also identified 8 distinct neuronal cell types that consist of both dopamine receptor expressing MSNs and inhibitory subtypes. These clusters were named based on their gene expression profiles and topography. Gene expression profiles were compared to previous datasets that profiled the NAc across mammalian species.

#### Label transfer between Xenium donors

A primary challenge when performing spatial transcriptomics on post-mortem human tissue is maintaining anatomical boundaries and landmarks during the freezing process. Flash freezing of fresh human tissue can result in neuroanatomical changes that can make brain region identification difficult. We identified that Br6660, one of our two brain donors, exhibited well-preserved neuroanatomy. Dimensionality reduction and clustering was performed within this donor and cell type labels were transferred to the second donor. To do this, *SpatialExperiment*^55^ objects were split into donor-specific objects, which were then converted to *Seurat*^61^ objects using the `as.Seurat()` function provided by *Seurat* v5.3.0^61^. Following conversion to *Seurat* objects, variable features were calculated using `FindVariableFeatures()` with default parameters and gene expression data was centered and scaled using `ScaleData()` with default parameters. Dimensionality reduction was performed within each donor. PCA was performed with `RunPCA()` with calculation of 50 principal components (PCs) and seed=2051. UMAP was used to further reduce dimensionality of the dataset with all 50 PCs calculated during the PCA step using the `RunUMAP()` function with `seed=2051`.

Following sample-specific dimensionality reduction in *Seurat*^61^ v5.3.0, label transfer was initiated with the `FindTransferAnchors()` function using 50 PCs and the reciprocal PCA algorithm. Br6660, the donor clustered with *BANKSY*, was used as the reference while data from Br6436 was used as the query. This function ultimately identifies related cells between the reference and query dataset in a reduced dimension space that serve as anchors for label transfer. Following identification of anchors, cell type labels were transferred using `TransferData()` using all 50 PCs. Cell type labels were then added to the *Seurat* object using `AddMetaData()`. Max prediction score was then plotted using *ggplot2* v3.5.2^62^.

### 4.7 Aligning Xenium tissues into a 3D common coordinate system

#### 3D reconstruction of NAc Xenium slices from a single donor

To reconstruct the Xenium sections in three dimensions, we used *Spateo*^63^ v1.1.1, a computational framework for multidimensional spatial transcriptomics analysis that supports scalable slice alignment and 3D reconstruction using a pairwise alignment pipeline. We used `st.align.morpho_align_transformation` to estimate pairwise transformations among adjacent sections of a single Xenium donor. Computation was performed on a GPU, and to improve scalability for Xenium-resolution data, we enabled `sparse_calculation_mode=True` and `use_chunk=True` with `chunk_capacity=2`, which follows *Spateo*’s sparse/chunked strategy for large spatial datasets. We also set `partial_robust_level=30` to allow robust partial matching between adjacent sections, since sections separated by 500 μm are not expected to exhibit complete spatial overlap, owing to biological structural changes across depth as well as sectioning-related deformation and edge truncation. This function returns the estimated transformation matrix, which is subsequently applied to the cellular spatial coordinates to generate aligned coordinates in the reconstructed reference frame.

After the initial global alignment, we performed a second refinement step for slice pairs that showed suboptimal alignment in the first pass, as determined by visual inspection and quantitative measures of cross-slice gene-expression similarity. For this refinement, we selected cell type subsets independently for each slice pair based on the clarity and anatomical informativeness of their spatial organization. Selected labels included astrocyte subtypes (Astro_A and Astro_B), excitatory neurons (Excitatory), choline acetyltransferase-positive neurons (CHAT), and segmented cells exhibiting both oligodendrocyte and medium spiny neuron characteristics (MSN_Oligo). We then re-estimated the transformation for these slice pairs based on cell type labels of subsetted cell populations using `st.align.morpho_align_transformation` with `rep_layer = [“CellType”]`, `rep_field = [“obs”]`, and `dissimilarity = [“label”]`. This refinement was intended to place greater emphasis on cell populations with stronger spatial structure, thereby reducing the influence of more heterogeneous or less anatomically informative cell classes during fine registration. Relative to the initial pass, we increased `partial_robust_level` from 30 to 50, 100, or 150 to improve robustness to partial overlap among these anatomically structured populations. We set `SVI_mode=False` so that the refinement was performed directly on the selected subset rather than using *Spateo*’s approximate large-scale inference mode. In this way, the first pass provided a coarse alignment of the full tissue sections, while the second pass improved local registration for poorly aligned slice pairs using biologically informative cellular landmarks.

#### Pairwise alignment evaluation based on cross-slice gene-expression similarity

We evaluated the quality of the 3D reconstruction using an established approach based on gene-expression similarity between each pair of adjacent aligned Xenium slices. The analysis was restricted to the intersection of the two sections’ spatial bounding boxes. At each tile size *h*, cells were assigned to matched square tiles, and tiles containing fewer than five cells in either section were excluded. A tile size was evaluated only when at least 10 matched tiles remained. For gene *g*, let 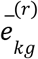 and 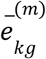 denote its mean expression in matched tile *k* of the reference (r) and moving (m) sections, respectively. These two spatial expression vectors are denoted as

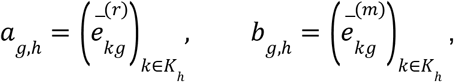

where *K*_*h*_ is the set of retained matched tiles. Similarity was calculated gene by gene across spatial tiles. We first calculated Pearson’s correlation coefficient (PCC) to measure linear agreement between the two spatial expression patterns as

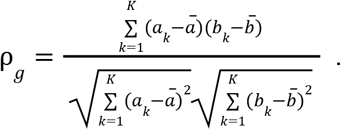

Mutual information (MI) was also used to capture more general, potentially nonlinear dependence:

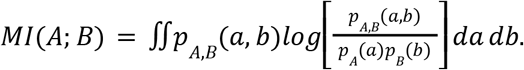

MI was estimated from the paired tile-level expression values using the *scikit-learn*^64^ v1.7.2 *k*-nearest-neighbor `mutual_info_regression` estimator with a fixed random seed. PCC ranges from ™1 to 1. MI is nonnegative and has no fixed upper bound. Higher PCC and MI values indicate greater cross-slice gene-expression similarity, corresponding to better alignment.

To capture agreement across spatial resolutions, scores were calculated at tile sizes of 200, 300, and 400 µ*m* and summarized by their median across scales. The final score for a section pair was the median across eligible genes.

### 4.8 Identifying changes along the anterior-posterior axis

#### 4.8.1 Changes in the cell composition

#### Grid-based cell type proportion analysis

To characterize variation in cellular composition throughout the reconstructed tissue volume, we performed a grid-based cell type proportion analysis using the aligned 3D coordinates from *Spateo*, corresponding to the medial–lateral (ML), dorsal–ventral (DV), and anterior–posterior (AP) anatomical axes. For each anatomical plane (ML–DV, ML–AP, and DV–AP), the projected coordinates were partitioned into a shared 5 × 5 lattice using equally spaced tile edges spanning the full aligned tissue volume. Cells within each two-dimensional tile were further stratified by position along the orthogonal anatomical axis: by tissue slice along the AP axis for the ML–DV plane and by 10 equally spaced intervals along the DV or ML axis for the ML–AP and DV–AP planes, respectively. Within each resulting spatial tile, the number of cells assigned to each cell type was divided by the total number of cells in that group to estimate local cell type proportions. This procedure enabled cellular composition to be compared across spatial positions along all three anatomical orientations.

#### 3D smoothing of cell type proportions

To estimate smooth spatial variation in cell type composition across the reconstructed tissue volume, we fit a binomial generalized additive model (GAM) to voxel-level cell type counts. This analysis was designed to complement the discrete grid-based summaries described above. Although the 5 × 5 grid analysis provides an interpretable local summary of cell type composition across AP depth, it produces many separate proportion profiles, which can be difficult to interpret as a global 3D pattern (**Fig. S6**). The GAM-based analysis therefore provides a model-based, smoothed representation of cell type proportion that can be visualized directly in three dimensions.

Using the *Spateo*-aligned in-plane coordinates and physical section depths, each cell *i* was represented by *S*_*i*_ = (*x*_*i*_, *y*_*i*_, *z*_*i*_), where *x*_*i*_ and *y*_*i*_ denote aligned spatial coordinates along ML and DV axes, respectively, and *z*_*i*_ denotes section depth along the AP axis. To reduce cell-level noise and obtain stable estimates of local cell type composition, the reconstructed tissue volume was discretized into spatial voxels. The aligned ML and DV coordinates were each divided into 50 quantile-based intervals, while the observed section depth defined the AP coordinate. This approach promoted more balanced cell occupancy along each in-plane axis and reduced sparsity relative to equal-width partitioning. Each voxel is defined by one ML coordinate interval, one DV coordinate interval, and one section depth.

For each voxel *v* and cell type *c*, we computed the total number of cells in the voxel, *N*_*v*_, and the number of cells assigned to cell type *c*, denoted *Y*_*v,c*_. The observed local cell type proportion was calculated as

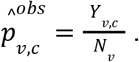

Voxels containing fewer than 20 total cells were excluded to avoid interpreting sparsely sampled tissue regions as representative of local cellular composition. For each retained cell type *c*, we modeled the voxel-level count as a binomial outcome:

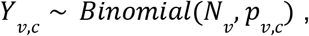

where *p*_*v,c*_ represents the underlying local proportion of cell type *c* in voxel *v*. The logit-transformed cell type proportion was modeled as a smooth function of the 3D voxel center:

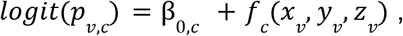

where (*x*_*v*_, *y*_*v*_, *z*_*v*_) denotes the voxel center and *f* (·) is a smooth spatial function. We represented *f*_c_ using a 3D tensor-product spline:

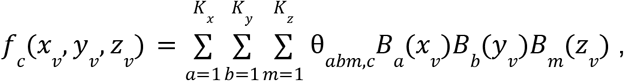

Where *B*_*a*_ (·), *B*_*b*_ (·), and *B*_*m*_ (·) are one-dimensional spline basis functions along the aligned *x, y*, and *z* directions, respectively. Their product forms a tensor-product basis function in 3D space, and θ_*abm,c*_ is the corresponding coefficient for cell type *c*. The spline coefficients were estimated by minimizing a penalized binomial negative log-likelihood:

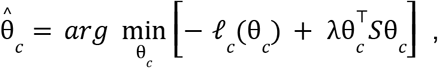

where *l*_*c*_ (θ_*c*_) is the binomial log-likelihood for cell type *c*, θ is the vector of spline coefficients, and *S* is the spline penalty matrix. The penalty term 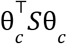 measures the roughness of the fitted tensor-product spline surface and discourages overly irregular spatial variation. The smoothing parameter λ controls the tradeoff between goodness of fit and smoothness. In this analysis, λ = 1. 0 was used to regularize the fitted spatial surface while retaining localized composition patterns. This model was fitted using *pyGAM* v0.12.0 with a logistic GAM and a tensor-product smooth term `te(0, 1, 2)`. We used *K*_*x*_ = 6, *K*_*y*_ = 6, and *K*_*z*_ = 4 spline bases for ML, DV, and AP orientations, respectively. To fit the grouped binomial model with *pyGAM* v0.12.0, each voxel was represented by two weighted binary observations: *Y*_*v,c*_ cells assigned to cell type *c* and *N*_*v*_ ™ *Y*_*v,c*_ cells assigned to all other cell types. After fitting the model, fitted cell type proportions were obtained at the observed voxel centers as

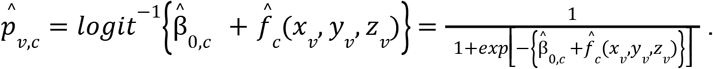

These fitted values provide smoothed estimates of local cell type proportion while borrowing information across nearby spatial locations.

To generate smooth 3D heatmaps, the fitted GAM for each cell type was evaluated at 60 × 60 × 20 regularly spaced prediction locations forming a three-dimensional lattice spanning the aligned ML, DV, and AP coordinate ranges. To restrict predictions to tissue-supported regions, we constructed a Delaunay triangulation of the occupied voxel centers using `scipy.spatial.Delaunay` and excluded prediction locations falling outside the triangulated volume using its `find_simplex()` method from *SciPy*^65^ v1.15.3. This procedure approximated the tissue volume by the convex hull of the occupied voxels and limited volume rendering to regions supported by observed data. For each cell type, the lower display threshold was defined as the 60^th^ percentile of the fitted proportions among locations retained by the tissue mask, such that volume rendering emphasized regions with relatively high fitted cell type proportions. This threshold was applied only during visualization and did not affect model fitting or prediction. This grouped-binomial framework accounts for differences in total cell numbers across voxels while allowing cell type proportions to vary smoothly across 3D tissue space.

#### 4.8.2 Changes in the gene expression

##### Spatial pseudobulk construction

Spatial gene-expression analysis was performed in the primary donor, Br6660, using the aligned Xenium coordinates from 11 sections spanning physical AP coordinates from 580 to 5580 µm. Increasing AP coordinate corresponds to the anterior-to-posterior direction. Raw counts were analyzed for a panel of 366 genes and 20 BANKSY-defined cell types. To compare corresponding absolute ML and DV locations across sections, the aligned ML–DV plane was divided into a common, globally anchored lattice of 2000 × 2000 µ*m* tiles. Let *S, k, c*, and *g* index physical slice, tile, cell type, and gene, respectively. For each slice–tile–cell type combination, raw counts for gene *g* were summed across its constituent cells to obtain the pseudobulk count *Y*_*Skcg*_. The corresponding exposure *A*_*Skc*_ was the sum of the morphology-derived nucleus-area size factors of those cells. Here, exposure denotes the standard count-model normalization quantity, representing the total nucleus-area-based normalization factor contributed by cells assigned to each pseudobulk.

Spatial support was determined independently of gene expression. For each cell type, eligible slices contained at least 100 cells and total exposure of at least 100. The selected AP interval was required to contain at least eight contiguous sections and span at least 60% of the complete AP range. Within this interval, a slice–tile pseudobulk required at least 10 cells and positive exposure, and a tile was retained only if it qualified in at least four sections. Gene–cell type combinations represented by fewer than two pseudobulks with nonzero counts or fewer than 10 transcripts in total were not tested. Selecting a contiguous AP interval and repeatedly observed tiles ensured that the AP coefficient was estimated from comparable tissue locations with adequate support across depth.

##### AP regression models

Each gene was modeled separately within each cell type using a log-link quasi-Poisson generalized linear model. Let *a*_*S*_ denote the physical AP coordinate in millimeters, centered within the selected cell type-specific AP interval. The primary model (M0) was

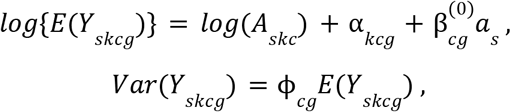

where α_*kcg*_ is a ML–DV tile fixed effect. Consequently, 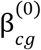 was estimated from within-tile differences across AP levels rather than from comparisons among different tissue locations. The coefficient represents the log change in normalized expression rate per 1-mm posterior displacement, conditional on a given tile. Effect sizes were reported as 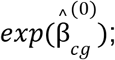 positive and negative values indicate increasing and decreasing expression rates, respectively.

To assess whether AP associations were sensitive to variation in local cell type density, we fitted a secondary model (M1) using the same pseudobulk rows as M0. A tissue-support mask was constructed from the cell coordinates and intersected with the tile to define the analyzed tissue area *T*_*Sk*_ for each section–tile combination. Cell type density was defined as *D*_*Skc*_ = *n*_*Skc*_/*T*_*Sk*_, where *n*_*Skc*_ is the cell count. Log density was decomposed using the analyzed-area-weighted mean for each slice into a within-slice component *W*_*Skc*_, and a between-slice component *B*_*Sc*_. The within-slice component represents local enrichment or depletion of a cell type in a tile relative to its slice-wide mean, whereas the between-slice component represents differences in the overall abundance of that cell type among sections. The sensitivity model was

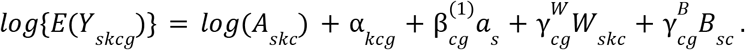

M1 was interpreted as a conditional sensitivity analysis because between-slice density could be correlated with AP position; M0 remained the primary model.

Models were fitted in R v4.5.0 using `glm` with `quasipoisson(link = “log”)`. Inference for the AP coefficient used a CR2 cluster-robust covariance estimator implemented with `vcovCR(type = “CR2”)` from *clubSandwich* v0.7.0, clustering by physical section. CR2 applies a bias-reduced small-sample correction to the cluster-robust covariance estimate to accommodate dependence among pseudobulk rows from the same section and prevent spatial tiles from being treated as independent AP replicates. AP coefficients were tested using `coef_test(test = “Satterthwaite”)` from *clubSandwich* v0.7.0. The Satterthwaite procedure assigns coefficient-specific approximate degrees of freedom according to the uncertainty in the robust variance estimate and provides a small-sample reference distribution when the number of independent section clusters is limited. Two-sided *p* values and 95% confidence intervals used these degrees of freedom. For the tested gene–cell type hypotheses, Bonferroni adjustment was applied separately to the M0 and M1 AP *p* values.

### 4.9 Characterizing cell type spatial relationships in Xenium data

#### Pairwise alignment of adjacent Xenium and Visium HD slices

The Visium HD windows had been placed to sample D1-island-rich regions across lateral, dorsal, ventral, and medial anatomical locations. This cross-platform alignment was performed to localize the Visium HD acquisition footprints within the reconstructed Xenium tissue and thereby refine the positions of D1_Island_A- and D1_Island_B-enriched pockets for the subsequent cell type spatial relationship analysis in the reconstructed Xenium 3D coordinate space. Each Visium HD section was paired with a Xenium section sampled at the nearest available physical depth. Eight pairs were analyzed, including six from donor Br6660 and two from donor Br6436, with separations of 20–50 µm between paired sections. The final within-donor Xenium coordinates obtained from the preceding Xenium alignment workflow were used as the fixed reference coordinates, and the corresponding Visium HD section was treated as the moving section.

The spatially annotated Visium HD objects, preprocessed and clustered following the steps described in Section 5.0, contained spatial coordinates expressed in full-resolution image pixels. For each sample, we obtained the sample-specific `microns_per_pixel` value from the Space Ranger `scalefactors_json.json` file and multiplied the pixel coordinates by this value to express them in micrometers. This conversion placed the Visium HD and Xenium spatial coordinates on the same physical scale. Before registration, the Visium HD *y* coordinate was reflected across the midpoint of its observed coordinate range to place the moving section in the same in-plane orientation as the fixed Xenium reference. Before feature-based alignment, Visium HD observations expressing fewer than 10 genes were removed with `scanpy.pp.filter_cells(min_genes=10)`, and genes detected in fewer than three Visium HD observations were removed with `scanpy.pp.filter_genes(min_cells=3)` from *Scanpy*^66^ v1.11.5. Xenium and Visium HD objects were then restricted to their common genes. The Xenium expression matrix was used in its precomputed normalized form. Visium HD counts were normalized using the library size calculated before gene intersection and were subsequently log-transformed. A joint principal-component representation of the fixed Xenium and moving Visium HD objects was computed with `align.group_pca()` from *Spateo*^63^ v1.1.1.

As with the within-donor Xenium alignment, cross-platform registration was performed using *Spateo*^63^ v1.1.1 in two stages: an initial alignment based on all observations that passed quality control, followed by label-guided refinement of pairs showing residual misalignment. Pairs requiring refinement were identified by visual inspection of the overlaid D1_Island_A, D1_Island_B, and white matter patterns. Because each Xenium–Visium HD pair was separated by only 20–50 µm, these shared anatomical structures within the Visium HD footprint were expected to exhibit nearly complete spatial overlap. We estimated the initial pairwise rigid alignment with `align.morpho_align_transformation()` using Xenium supplied as the fixed model and Visium HD as the moving model. The joint PCA coordinates were provided through `rep_layer = “X_pca”` and `rep_field =“obsm”`, and cross-modality feature dissimilarity was calculated using cosine distance with `dissimilarity = “cos”`. Alignment was performed on GPU with `max_iter = 500` and `partial_robust_level = 100`. To support the size of the spatial datasets, we enabled `sparse_calculation_mode=True` and `use_chunk=True` with `chunk_capacity=2`. The fitted transformation was applied with `align.morpho_align_apply_transformation()` to the complete, unfiltered Visium HD object to yield aligned coordinates for every Visium HD observation in the reconstructed Xenium 3D coordinate space.

For pairs requiring further refinement, a second rigid transformation was estimated directly in the first-pass aligned coordinate space. Visium HD observations were subset to anatomically structured labels shared across the two modalities, including D1_Island_A, D1_Island_B, and a harmonized CHAT_PVALB_Inh category, where Xenium CHAT and Inh_PVALB cells were mapped to the Visium HD CHAT_PVALB spatial domain. The residual transformation was fitted with `align.morpho_align_transformation ()` using `rep_layer = [“CellType”]`, `rep_field = [“obs”]`, and `dissimilarity = [“label”]`. We set `SVI_mode = False` so that fitting was performed directly on the selected observations. The resulting residual transformation was then applied with `align.morpho_align_apply_transformation ()` to all first-pass Visium HD coordinates. One pair of Xenium and Visium HD sections required iterative white-matter-guided refinement. The first iteration restricted fitting to white-matter observations within the tissue-wide overlap supported by cross-modality nearest-neighbor matches within 100-µm, and the second iteration used WM-to-WM matches within 300 µm in the updated coordinate space. Nearest-neighbor searches in both iterations were performed using `spatial.cKDTree.query()` from *SciPy*^65^ v1.15.3. Alignment quality was assessed by visual inspection of pairwise Xenium–Visium HD overlays for shared anatomical labels of D1_Island_A, D1_Island_B, and white matter (**Fig. S14**).

#### Characterizing cell type spatial relationships across ROIs and AP depth

We used Cell type Relationship Analysis Workflow Done Across Distances (*CRAWDAD*^31^) to characterize how directional cell type spatial relationships varied among anatomically defined regions of interest (ROIs) and across the AP axis of the reconstructed nucleus accumbens. *CRAWDAD* compares the observed cell type composition surrounding a reference cell population with empirical null distributions generated by locally shuffling cell type labels at a series of spatial length scales, thereby distinguishing spatial enrichment from depletion while preserving tissue-scale variation in cell type composition.

ROIs were defined in the aligned Xenium coordinate system using the six Visium HD acquisition footprints described in the preceding cross-platform alignment analysis. All polygon construction, combination, and spatial-assignment operations were performed using *Shapely*^67^ v2.1.1. For each Visium HD window, its aligned observation coordinates were converted to a polygon using `concave_hull()` with `ratio = 0.05` and `allow_holes = False`; the minimum rotated rectangle of this concave hull was then used as the window footprint. One Visium HD window had been positioned to investigate D1_Island_A and D1_Island_B populations in the lateral nucleus accumbens and was therefore used to define the lateral ROI. The other five footprints were combined using `unary_union()` to define the medial region. Areas in which the lateral and medial footprints overlapped were defined as lateral. The medial region was subdivided into dorsomedial and ventromedial ROIs using a single dividing line in the aligned dorsal–ventral coordinate. To reduce sensitivity to extreme coordinates, the line was defined as the midpoint between the 0.05th and 99.95th percentiles of the aligned *y* coordinates of broad medium spiny neuron populations located within the medial region. These populations comprised DRD1_MSN, DRD2_MSN, D1_Island_A, D1_Island_B, and MSN_Oligo. Medial cells with coordinates greater than or equal to this midpoint were assigned to the dorsomedial ROI, whereas cells below the midpoint were assigned to the ventromedial ROI. Xenium cells were assigned to the resulting mutually exclusive polygons using `intersects_xy()`; cells outside all three primary polygons were labeled outside_global_roi.

*CRAWDAD*^68^ v1.0.1 was applied independently to each Xenium section and analysis region using the final aligned *x* and *y* coordinates and the previously assigned cell type labels. Each section–region combination constituted a separate two-dimensional analysis task. We used a fixed universe of 20 cell types, and a cell type was eligible to serve as a reference population in a given task only when at least 20 cells of that type were present. Cell coordinates and annotations were converted to a spatial feature object using `toSF()`.

We used a neighborhood distance of *d* = 50 µ*m*, with overlapping neighborhoods merged and duplicate neighboring cells removed by setting `removeDups = TRUE`. The reference cells used to seed each neighborhood were excluded from the neighbor-composition calculation. Null datasets were generated with `makeShuffledCells()` using square-tile shuffle scales from 200 to 1000 µ*m* at 100 µ*m* intervals. We generated three permutations per scale using `perms = 3` and `seed = 1`. Directional spatial relationships were estimated using `findTrends()` with `neighDist = 50` and the scale-specific shuffled datasets. A directional reference–neighbor relationship describes whether a candidate neighbor cell type is enriched or depleted within the neighborhoods surrounding cells of the reference type. Results were converted to long format with `meltResultsList(withPerms = TRUE)`. The three permutation-specific *Z*-scores were averaged for each section, ROI, directional cell type pair, and shuffle scale. Statistical significance was evaluated using a single two-sided Bonferroni-corrected threshold of |*Z*| ≥ 3. 84, based on the fixed universe of 20 cell types and 400 directional reference–neighbor tests. Positive values meeting this threshold were classified as significantly enriched, and negative values meeting it were classified as significantly depleted. For each eligible directional pair, the first significant scale was defined as the smallest evaluated shuffle scale at which the absolute mean *Z*-score reached 3.84. Spatial relationships across AP depth were compared among sections ordered by physical AP coordinate, while regional differences were compared among the lateral, dorsomedial, ventromedial, and outside_global_roi compartments. All comparisons used the same *CRAWDAD* parameters and significance threshold.

### 5.0 Visium HD Data Analysis

#### 5.0.1 Raw Data Processing

Raw imaging-based sequencing data was preprocessed with *SpaceRanger* v4.0.1 using the `spaceranger count` function within the *SpaceRanger* v4.0.1 software with the GRCh38 human genome. *SpaceRanger* v4.0.1 software also provides segmented output based on *StarDist*^69^. To achieve single cell resolution spatial transcriptomics, segmented outputs were loaded into R v4.5.0 using `read10xCounts()` from *DropletUtils* v1.28.0 on a per-sample basis. Cellular spatial coordinates were loaded via the `cell_segmentations.geojson` file output from *spaceranger count*. Polygons from the `cell_segmentations.geojson` file were converted to centroids with the `st_centroid()` from the *sf*^70,71^ package version 1.0-21. Centroids were then converted into a plain matrix with st_coordinates() from the *sf*^70,71^ package version 1.0-21. *SpatialExperiment* objects were generated using *SpatialExperiment* v1.18.1 and then converted to a *SpatialFeatureExperiment*^72^ containing cell polygons with *SpatialFeatureExperiment* v1.10.1. Finally, per-sample *SpatialFeatureExperiment* objects were combined with `cbind()`.

#### 5.0.2 Quality Control

Following the generation of a merged *SpatialFeatureExperiment* object, we next calculated the number of genes, unique molecular identifiers (UMIs), and percentage of reads mapping to the mitochondrial genome on a per-segmented cell basis. The use of segmented cell outputs from *SpaceRanger* allowed us to effectively treat the data as single nucleus RNA-sequencing data. Therefore, we calculated adaptive thresholds on a per-sample basis to identify uninformative cells based on low number of genes, low number of UMIs, and high percentage of reads mapping to the mitochondrial genome using the `isOutlier()` function from *scuttle*^56^ v1.18.0. Uninformative cells were defined as those with cells with a number of genes and number of UMIs less than 2.5 median absolute deviations (MADs) from the sample median, and a percentage of reads mapping to the mitochondrial genome greater than 5 MADs from the sample median. VisiumHD sample preparation requires direct tissue targeting with a cassette. During this process, the cassette can be moved on the tissue sample, resulting in the unexpected inclusion of cells, which are not fully profiled. One sample, H1-XNQ4F2B_A1, contained extraneous cells that were removed based on their spatial coordinates. Following removal of uninformative cells, our dataset consisted of 439,451 segmented nuclei across 8 samples.

#### 5.0.3 Spatial Clustering with BANKSY

A primary goal of the VisiumHD data was to understand whether transcriptionally distinct cell types form topographically organized spatial domains. Thus, we performed spatial clustering with *BANKSY*^60^ v1.4.0. To reduce the computational resources and time required to run this analysis, we identified the top 2000 highly variable genes by running `modelGeneVar()` followed by `getTopHVGs()` from *scran*^73^ v1.36.0 on the entire pooled dataset. Gene selection was insensitive to blocking by `sample_id`, or array, with 1,931/2,000 genes shared with a blocked fit. The object was subsetted to the top 2000 highly variable genes and the augmented BANKSY matrix was calculated. Specifically, samples were again processed individually for `computeBanksy()` with `compute_AGF = TRUE`. The azimuthal gabor filter, or AGF, detects gene expression gradients which are important for understanding topographical domains within the human NAc. The BANKSY augmented matrix was calculated using `k_geom = c(25,50)`, resulting in the incorporation of information from the nearest 25 and 50 neighbors. The use of two `k_geom` values allows for the identification of both fine-grained local neighbors and broad spatial domains. Following calculation of the BANKSY augmented matrix, principal component analysis (PCA) was performed with `runBanksyPCA()` with “Sample” as the grouping factor, `seed=1000`, and `use_AGF=TRUE`. Batch correction was not used as it could have erased any biological heterogeneity that exists along different anatomical axes given that our Visium HD arrays were placed along different aspects of the AP/DV/ML axes. For spatial domains, we chose to set ***λ***=0.8, allowing for the inclusion of nearest neighbors and spatial information. Uniform manifold approximation and projection (UMAP) was used to further reduce the dimensionality of the data for plotting purposes. UMAP was generated using `runBanksyUMAP()` with ***λ***=0.8 and `seed=1000`. Following dimensionality reduction, clustering was performed with the `clusterBanksy()` function with `use_agf=TRUE`, ***λ***=0.8, `seed=948`, and a resolution of 0.4. For Visium HD data, we used the Leiden algorithm for clustering, resulting in 12 transcriptionally distinct spatial domains.

#### 5.0.4 Label Transfer

To understand the cell type composition of spatial domains within the Visium HD data, we used label transfer to predict the transcriptionally distinct cell type each segmented cell represents. Using a recently published snRNA-seq atlas of the human NAc from our lab, we performed label transfer using a combination of algorithms and strategy. First, segmented cell identify was predicted using *RCTD* (Robust Cell Type Decomposition)^74^, a supervised learning method that was originally built to identify single cells in spot-based spatial transcriptomics technologies where several cells can be captured in a single spots, which is implemented within the *spacexr*^74^ package v1.0.0. RCTD also provides several “modes” which work under the assumption that each segmented cell may contain 1 or many cell types. For this dataset, RCTD was performed using doublet mode which constrains the model to predict at most 2 cell types for each segmented cell, with a minimum UMI value of 20. RCTD provides a class for each prediction with classes of singlet, doublet_certain, doublet_uncertain, or reject. >75% of cells within the Visium HD dataset were classified as either singlet, doublet_certain, or doublet_uncertain (**Fig. S10A**). Cells labeled as putative doublets likely arise from transcript leakage from neighboring astrocytes or oligos, or incomplete segmentation resulting in transcripts from non-neuronal cell types being included in the neuronal signature. This is evident as a large percentage of cells primarily labeled neurons are secondarily classified as astrocytes or glia (**Fig. S10B**). To recover the cells RCTD was not confident in providing labels, we next relied on SingleR, a computational method that predicts cell types based on correlation of marker genes between the two datasets, which is implemented in the *SingleR*^75^ package v2.10.0. SingleR provides “pruned.labels”, or high-confidence calls, which were used to label any remaining cells not provided a cell type by RCTD. Of the 439,451 segmented cells, only 6 were not successfully classified. 3 of these cells had UMI counts of 1 for every detected gene resulting in a failure to fit the RCTD model. The final 3 cells were labeled as rejected by RCTD and had no pruned label. Due to the failure of two separate algorithms to predict a cell type, these cells were dropped from further analysis.

#### 5.0.5 Identifying differences in cellular composition with crumblr

To understand whether D1-island spatial domains exhibited differential enrichment of transcriptionally distinct D1-MSN subtypes, we next performed a cellular composition analysis using *crumblr*^25^ v1.0.0. Raw counts from VisiumHD data were pseudobulked using the `aggregateToPseudobulk()` function provided by *dreamlet*^76^ v1.9.2 with `cluster_id = “snRNA_label”` and `sample_id = “Sample”`. Sample was defined as a combination of Visium HD array and spatial domain resulting in per-array, per-spatial domain counts for each of the D1-MSN subtypes. Cell counts were then extracted with `cellCounts()` and normalized with a centered log-ratio transformation, via the `crumblr()` function, to convert compositional counts to log-ratio values that can be used for linear modeling. Differential cellular composition was then tested using a linear mixed model fit with the `dream()` function from the *variancePartition* package^76–78^ v1.40.2 using the formula `~ 0 + ‘Spatial_Domain’ + (1 | sample_id)`. This formula yields one coefficient per spatial domain and the random intercept absorbs array-specific baseline shifts in cellular composition, allowing for within array testing. Differences between D1-island spatial domains were identified by defining the contrast D1_Island_A - D1_Island_B with `makeContrastsDream()` function from the *variancePartition* package^76–78^ v1.40.2. `eBayes()` from the *limma*^79^ package v3.64.3 was then used to perform statistical testing.

### 5.1 Cross-species comparisons of medium spiny neurons

#### MetaNeighbor

MetaNeighbor^29^ was used to understand the molecular conservation of medium spiny neurons across technologies and species. Metaneighbor is a computational algorithm that generates cell-to-cell networks based on shared transcriptional signatures. Conservation between datasets or species is measured as the ability for a cell type to identify the corresponding cell types across the similarity networks and quantified with area under the receiver operator characteristic (AUROC). For this analysis, we included MSN containing cell types and domains from human Xenium, human snRNA-seq, human Visium HD, NHP snRNA-seq, and rat snRNA-seq. These datasets span different datasets and technologies which require the identification of a unified gene set consisting of 1-to-1 orthologs across species. To identify the unified gene set, NHP and rat genes were first converted to 1-to-1 orthologs using the `convert_orthologs()` function from the *orthogene*^80^ package v1.14.0 with the *gprofiler* method (`method=“gprofiler”`). Any gene representing a 1-to-many or many-to-many orthologs were dropped from further analysis (`non121_strategy = “drop_both_species”`). The unified gene set was finally generated using `intersect()` from *dplyr*^81^ v1.1.4 and included 295 genes. Each R object specific to each dataset included in this analysis was then subsetted to include only genes within the unified gene set of 1-to-1 orthologs. Following the generation of a single object containing all datasets with `cbind()`, the unsupervised low-memory version of MetaNeighbor was applied with `MetaNeighborUS(fast_version = TRUE)` within the *MetaNeighbor* package v1.28.0 with each cell type tested against each other (`one_vs_best = FALSE`). Importantly, we chose not to restrict the analysis to only highly variable genes and instead used all genes within the unified gene set (`var_genes` `= rownames(combo)`). AUROC values between cell types were then plotted with `Heatmap()` from the *ComplexHeatmap*^82^ package v2.24.0 with the order of rows and columns defined by the dendrogram generated from the `plotHeatmap()` function included in the *MetaNeighbor*^29^ package v1.28.0.

#### Correlation of transcriptional signatures between human and NHP islands

To investigate the conservation of marker genes between human and NHP islands, we first calculated marker genes within each dataset separately. Specifically, we performed one-versus-all testing on the log-normalized count matrices with the `findMarkers_1vALL()` function provided by the *DeconvoBuddies*^83,84^ package v1.0.0 with sample as the blocking term to account for any differences in transcript abundance between samples. Following calculation of marker genes for both datasets, only genes with a 1-to-1 ortholog were kept for further analysis. 1-to-1 orthologs were identified with the `convert_orthologs()` function from the *orthogene*^80^ package v1.14.0 with the “gprofiler” method (`method = “gprofiler”`). The standardized log-fold change (`std.logFC`) for 1-to-1 orthologs were then plotted with *ggplot2*^62^ v3.5.2.

### 5.2 RNAScope

#### 5.1.1 Single-molecule fluorescence *in situ* hybridization (smFISH)

Six 10 µm tissue sections from donor Br6660 were selected at different AP levels (anterior, intermediate, and posterior, 1000µm apart) for single molecule fluorescence in situ hybridization (smFISH) using the RNAscope Multiplex Fluorescent Reagent Kit v2 (Cat #323110, ACD, Hayward, California) as previously described^18^. Slides were fixed in Neutral Buffered Formalin Solution (NBF) (Ct # HT501128-4L, Sigma-Aldrich, St. Louis, Missouri) for 30 min at room temperature. A series of dehydration steps followed by a 10 min pretreatment with hydrogen peroxide and a 30 min incubation with Protease IV at room temperature were performed. Two different probe panel combinations were used to delineate the two distinct D1-island subtypes. Panel one included: Hs-VIP (Cat No. 452751, ACD, Hayward, California), Hs-RXFP1 (Cat No. 422821-C2, ACD, Hayward, California), Hs-DRD1 (Cat No. 524991-C3, ACD, Hayward, California), Hs-MBP (Cat #411051-C4, ACD, Hayward, California). Panel 2 included: Hs-MBP (Cat #411051-C1, ACD, Hayward, California), Hs-SEMA5B (Cat No. 52299-C2, ACD, Hayward, California), Hs-PROK2 (Cat No. 437331-C3, ACD, Hayward, California), Hs-DRD1 Cat No. 524991-C4, ACD, Hayward, California). Tissue sections were stored at 4 °C in 4 x SSC buffer overnight, followed by probe labeling. After amplification steps (AMP 1-3), the probes were fluorescently labeled using the following Opal Dyes at 1:500 concentration (Perkin Elmer, Waltham, MA; diluted 1:500) combination per panel: Opal 570 for Channel 1 (Hs-VIP), Opal 620 for Channel 2 (Hs-RXFP), Opal 690 for Channel 3 (Hs-DRD1), and Opal 520 for Channel 4 (Hs-MBP), Opal 520 for Channel 1 (Hs-MBP), Opal 570 for Channel 2 (Hs-SEMA5B), Opal 620 for Channel 3 (Hs-PROK2), and Opal 690 for Channel 4 (Hs-DRD1). Sections were counterstained with DAPI (ACD, Hayward, California) to mark nuclei and coverslipped with

Fluoromount-G. Images were taken using the Akoya PhenoImager Fusion slide scanner with *Fusion* software (v2.3.1) at 20x magnification using a 10x objective. The resulting widefield epifluorescence images were spectrally unmixed using InForm Software (v3.1.0) to remove lipofuscin autofluorescence.

### 5.3 Non-negative matrix factorization of rat snRNA-seq data

#### Cross validation to determine optimal rank and NMF model fitting

Non-negative matrix factorization^26^ is a dimensionality reduction technique that decomposes a large matrix into two smaller matrices, termed W and H, such that:

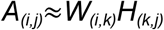

Where *i* represents all genes within the snRNA-seq dataset, *j* represents all cells within the snRNA-seq dataset, and *k* represents the optimal rank, or number of factors. *W* is a non-negative matrix consisting of genes as rows and the identified latent factors as columns. *H* is a non-negative matrix where latent factors are rows and cells within the dataset are columns. Non-negative matrix factorization is particularly suited to transcriptomics data as transcript counts are >0 and do not contain any negative values. The optimal rank (*k*) was determined using the log-normalized counts matrix from the rat snRNA-seq dataset with `cross_validate_nmf()` from *singlet*^85^ v0.99.6 with ranks of 30, 40, 50, 60, 70, 80, 90, 100, and 125 tested. For cross validation, 3 random initializations (`n_replicates = 3`) were used to generate a stable error estimate across a maximum of 100 iterations (`maxit = 100`) with an L1/LASSO penalty of 0.1 (`L1=0.1`) and an L2/ridge-like penalty of 0 (`L2=0`). Additionally, 20% of the gene expression matrix was withheld as a test set during each model fitting. Cross validation identified an optimal rank of 53 NMF patterns (**Fig. SXXX**).

Following the identification of the optimal rank with cross validation, a final NMF model fitting was performed with an optimal rank of 53. Specifically, NMF was implemented with the `nmf()` function from the RcppML^86^ package v0.5.6 with an L1/LASSO penalty of 0.1 (`L1=0.1)`, a convergence tolerance of 1×10^−6^ (`tol = 1e-06`), and a maximum iteration of 1000 (`maxit = 1000`). Non-negativity constraint was ensured for both the W and H matrices (`nonneg=TRUE`) and a scaling diagonal was used such that the columns and rows within w and h sum to 1 (`diag = TRUE`). Finally, zeros within the matrix were treated as observed zero values and not missing data (`mask_zeros = FALSE`).

#### Projection of NMF patterns into target datasets with transfer learning

To identify shared transcriptional signatures across species with NMF, the 53 patterns identified within the rat were then projected into a mouse Xenium and human Visium HD dataset using the `project()` function within the RcppML^86^ package v0.5.6. To avoid additional sparsity within the target dataset the L1/LASSO penalty was set to 0 in the `project()` function (`L1=0`). The target datasets represent a new dataset (*A’*) consisting of genes as rows and cells as columns. The *W*, matrix consisting of genes as rows and NMF patterns as column, stays constant during projection such that we are solving for *H’*, or a new matrix consisting of NMF patterns as rows and cells within the target dataset as column:

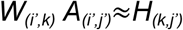

Importantly, it represents genes found within both the reference and target dataset. Before projection, the W matrix is subsetted for only 1-to-1 orthologs. These orthologs were identified with the `convert_orthologs()` function from the *orthogene*^80^ package v1.14.0 using the *gprofiler* method (`method=“gprofiler”`). Any gene representing a 1-to-many or many-to-many orthologs were dropped from further analysis (`non121_strategy = “drop_both_species”`). Following conversion of the target dataset genes to 1-to-1 orthologs, the common genes between the reference and target dataset were identified using `intersect()` from *dplyr*^81^ v1.1.4. For the human Visium HD dataset, projection was completed on a per-sample basis with cell-level loadings normalized per-factor to a sum of 1 across all cells within each sample. For the mouse xenium dataset, projection was completed on the entire dataset with cell-level loading normalized per-factor to a sum of 1 across all cells in the dataset. Normalized cell-level loadings were appended to the metadata or column data of the associated object and used for visualization.

### 5.4 Assessing genetic risk with scDRS

Disease-associated gene sets were derived from LDSC-munged GWAS summary statistics using *MAGMA*^87^ v1.10 and single nucleotide polymorphism (SNP) location files and linkage disequilibrium reference panel was derived from 1000 Genomes european ancestry phase 3 reference^88^. Specifically, we used the default single nucleotide polymorphism (SNP)-wise mean model which aggregates SNP *p*-values into a single gene-wide *p*-value determining whether the gene is associated with the disorder. We also annotated all SNPs within 35 kilobases (kb) upstream of the gene start and 10 kilobases (kb) downstream of the gene end. These values were previously used to identify cell types important for Parkinson’s disease^89^ and the expanded window also aids in capture of distal enhancer and promoter elements in which variation could affect gene expression. Importantly, gene coordinates were reported in hg19 and restricted to only the genes present in the VisiumHd dataset. Disease-associated gene sets contained the 1,000 genes with the highest gene-level Z-statistic per trait and weighting each gene by that statistic. The LDSC-munged GWAS summary statistics do not contain a p-value, and thus two-sided-p-values were derived under a standard normal null distribution using the formula *p*=2ϕ(−|Z|). MAGMA requires unique SNPs and thus SNP identifiers were de-duplicated keeping the first instance of the SNP. Duplicates ranged in 6-20 per trait, except for GSCAN traits which contained ~990 duplicated SNPs. The VisiumHD dataset was subsetted to exclude any excitatory cells or those derived from the Hypo spatial domain as they are not within the NAc. Following the subset, 407,397 cells were normalized to 10,000 counts per cell and log1p-transformed with the intercept, log number of detected genes, and sample indicators were regressed out. Genes were then binned into a 20 × 20 grid by mean expression and variance, which was then used to match control gene sets. Per-cell disease scores were then calculated using *scDRS*^32^ v1.0.2 against 1,000 expression-matched control gene-sets. Group-level association was tested by comparing the 95th percentile of normalized scores within each spatial domain or cell type against the 95th percentile of normalized scores from each of the 1,000 expression-matched control gene sets. This test generates a Monte-Carlo *p*-value and z-score that was then corrected for multiple testing across disease/trait x spatial domain/cell type using the Benjamini-Hochberg^90^ correction.

### 5.5 Identifying druggable cell type targets with drug2cell

drug2cell^52^ is an open-source software that tests whether known drug targets are enriched within single cells. Known drug targets are taken from ChEMBL, which is a database containing information regarding small molecules, drug targets, and mechanism of action, within the Phase 4-restricted dictionary. We augmented the drug dictionary to include the psychedelic drugs psilocybin, psilocin, lysergic acid diethylamide (LSD), mesacline, and N,N-demethyltryptamine (DMT). Per-cell scores were computed with `drug2cell.score(method=“mean”)` on log-normalized counts of the same 407,397 cells included in the scDRS analysis. Drugs enriched in each spatial domain were identified using the Wilcoxon rank-sum^91^ tests of each group against all remaining cells with Benjamini-Hochberg^90^ adjusted *p*-values.

### 5.6 Acknowledgements, Funding, Authorship Contributions

All Xenium and VisiumHD spatial transcriptomics data have been deposited at the Neuroscience Multi-Omic Data Archive (NEMO) under accession nemo:col-s93bogv. All original code has been deposited at https://github.com/LieberInstitute/xenium_NAC and is publicly available at 10.5281/zenodo.22755736 as of the date of this publication. Additionally, we provide interactive web-based app for exploration of VisiumHD and Xenium data that are linked within the GitHub README page. Any additional information required to reanalyze the data reported in this paper is available from the lead contacts upon request.

### 5.7 Acknowledgements, Funding, Authorship Contributions

## Acknowledgements

We gratefully thank the families who donated tissue to make this research possible. We thank the Essel Foundation and the families of Connie and Stephen Lieber and Milton and Tamar Maltz for their generous support of this work. We would like to acknowledge the contributions of Dr. Fernando Goes, the late Dr. Llewellyn B. Bigelow, Amy Deep-Soboslay, Anna Brandtjen, and James Tooke for sample curation and clinical characterization. We also thank the Office of the Chief Medical Examiner of the State of Maryland for their collaboration on tissue collection. We thank members of the Martinowich, Maynard, Page, and Hicks labs for critical reading of the manuscript and feedback. We thank Dr. Jeremy Day and his lab members for generously sharing their data ahead of print. We thank the Joint High Performance Computing Exchange (JHPCE) for providing computing resources for these analyses. The Xenium Analyzer runs were conducted at the Experimental Computational Genomics Core, Johns Hopkins School of Medicine. Portions of some figures were created with BioRender.com.

## Author Contributions

Conceived and Designed the Study: KM, KRM, SCH

Performed Experiments and Collected Data: SVB, IDRA, YD, SEM, RZ Software: RAP, JY, RAM

Formal Analysis: RAP, JY Data Curation: RAP, JY, RAM Tissue Resources: JEK, TMH

Writing – original draft: RAP, JY

Writing – review & editing: RAP, JY, SVB, KM, KRM, SCH Visualization: RAP, JY, SVB, IDR

Supervision: KM, KRM, SCH Funding acquisition: KM, KRM, SCH

Project Administration: KM, KRM, SCH

## Supplementary tables

**Table S1**. Xenium panel consisting of 366 total genes.

**Table S2**. Marker gene statistics from 1-versus-all testing between Visium HD spatial domains. All gene information is human-specific.

**Table S3**. Marker gene statistics from 1-versus-all testing between NHP MSN types. All gene information is NHP-specific.

## Supplementary Videos

**Video S1**. 3D rendering of D1-island subtypes identified with Xenium spatial transcriptomics.

**Video S2**. 3D smoothed surfaces of D1_Island_A. **Video S3**. 3D smoothed surfaces of D1_Island_B. **Video S4**. 3D smoothed surfaces of White Matter. **Video S5**. 3D smoothed surfaces of Excitatory.

