## Supplementary Figures for "Three dimensional reconstruction of the human nucleus accumbens reveals topographic organization and molecular heterogeneity of D1-islands across the anterior posterior axis"

### Supplemental Figures

**Figure S1**. **Donor Br6660 frozen coronal slab images. A.** The anterior face of the coronal slab at the level of anterior striatum (left) and corresponding schematic with labeled structures (right). CC - corpus callosum, CN - caudate nucleus, IC - internal capsule, Pu - putamen, VS - ventral striatum, WM - white matter. **B.** The posterior face of the same coronal slab as in A at the level of the ventral pallidum (left) and corresponding schematic with labeled structures (right). AC - anterior commissure, GP - globus pallidus, Hyp - hypothalamus, OT - optic tract, VP - ventral pallidum. **C.** Br6660 brain block images at different depths along the AP axis 500 - 1000 μm apart with labeled structures. Purple boxes represent levels at which sections were processed for Xenium assay.

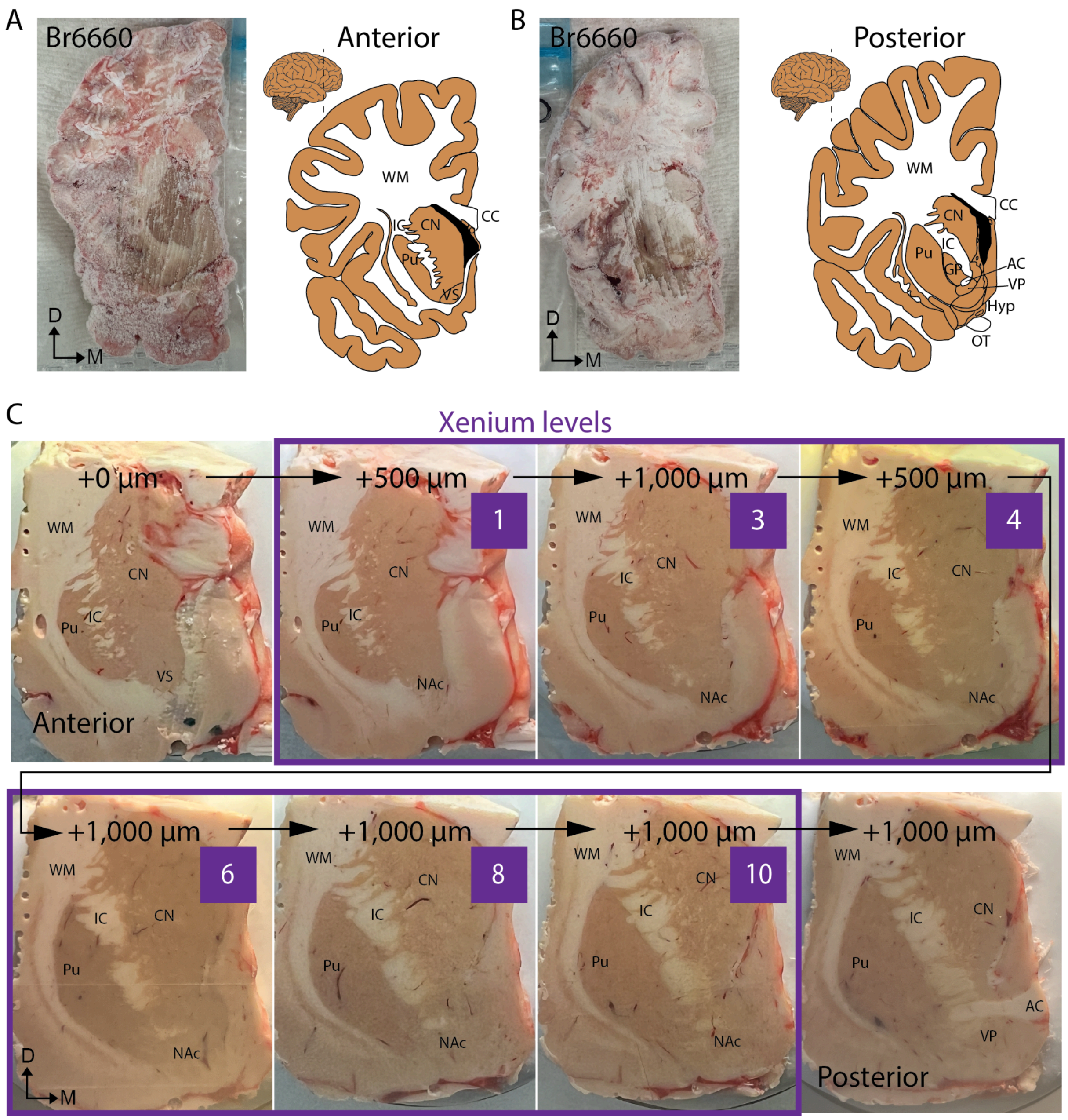

**Figure S2**. **Donor Br6436 frozen coronal slab images. A.** The anterior face of the coronal slab at the level of anterior striatum (left) and corresponding schematic with labeled structures (right). LV - lateral ventricle, St - striatum, WM - white matter. **B.** The posterior face of the same coronal slab as in A at the level of ventral pallidum (left) and corresponding schematic with labeled structures (right). AC - anterior commissure, GP - globus pallidus, Hyp - hypothalamus, OT - optic tract, VP - ventral pallidum. **C.** Br6436 brain block images at different depths along the AP axis 500 - 1000 µm apart with labeled structures. Purple boxes represent levels at which sections were processed for Xenium assay.

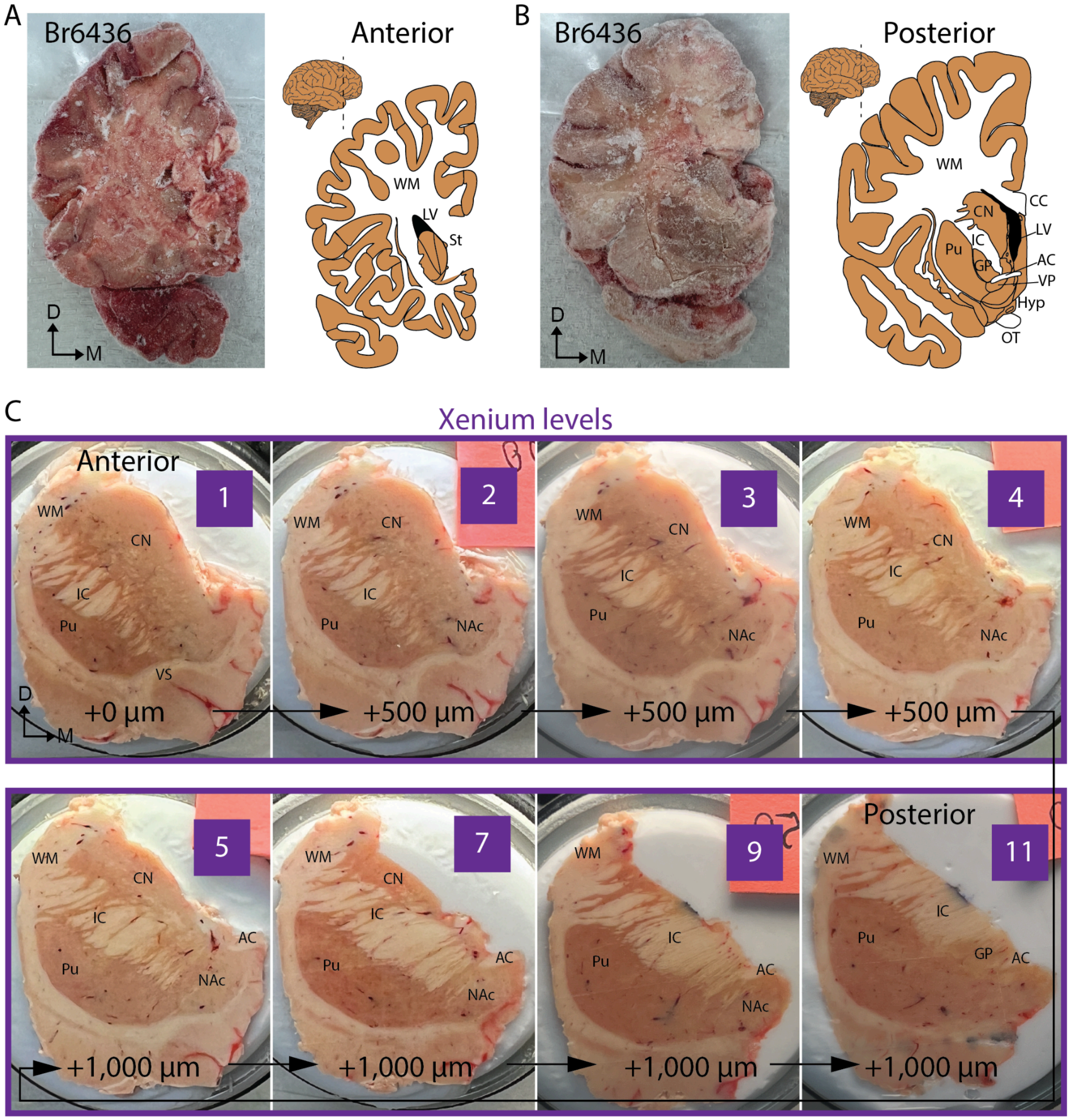

**Figure S3**. **Quality control of Xenium cells across tissue sections. A.** Distributions of the four cell-level QC metrics: total gene-expression transcripts, number of detected genes, percentage of negative-control and unassigned transcripts, and cell area. **B.** Number of outlier cells identified by each QC criterion and the percentage of cells removed from each section after applying all criteria.

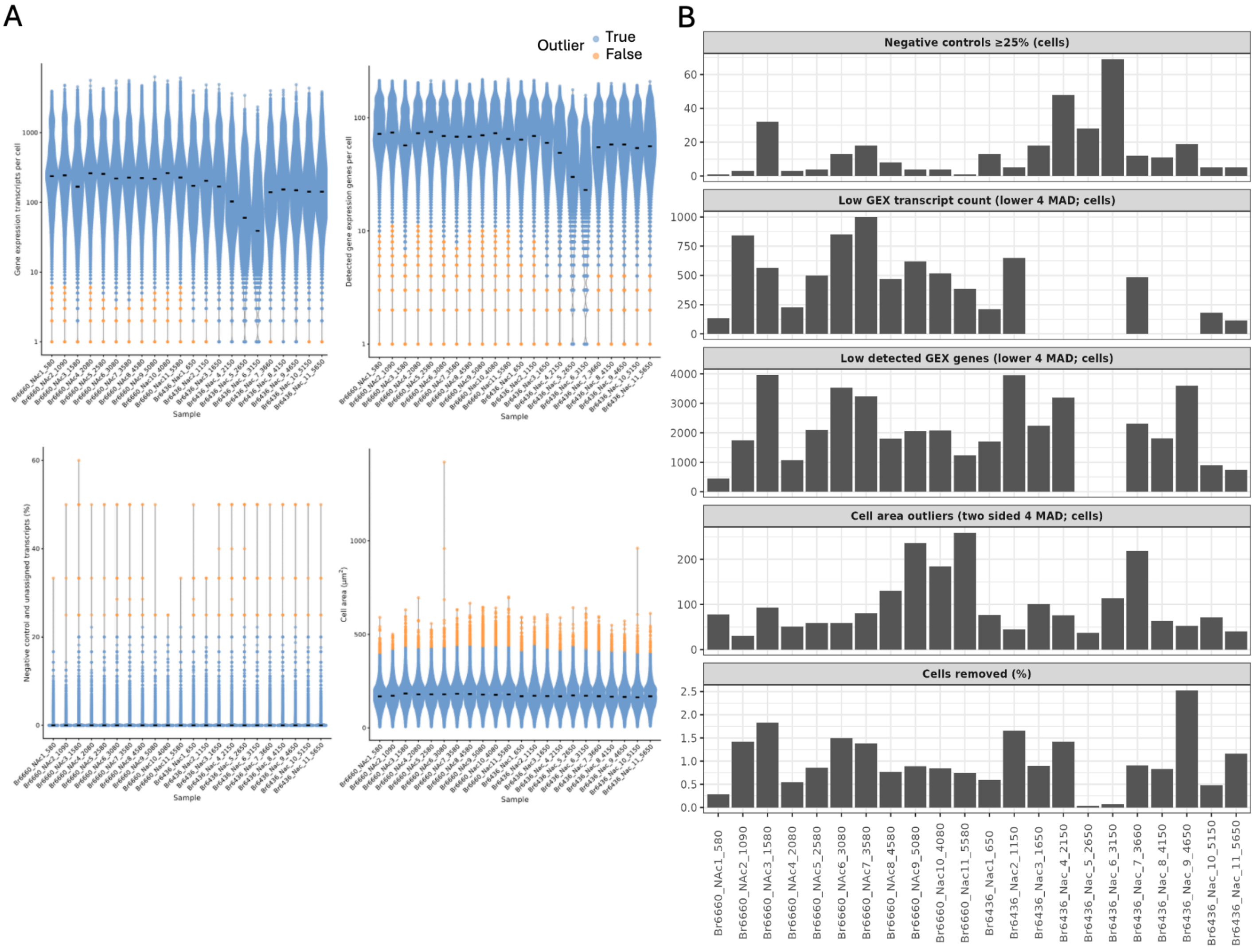

**Figure S4**. **Spatial registration between Xenium cell types and previously reported spatial domains and cell types**. Heatmaps depicting spatial registration results between **A**. Xenium and Visium spatial domains, and **B.** Xenium cell types and snRNA-seq cell types.

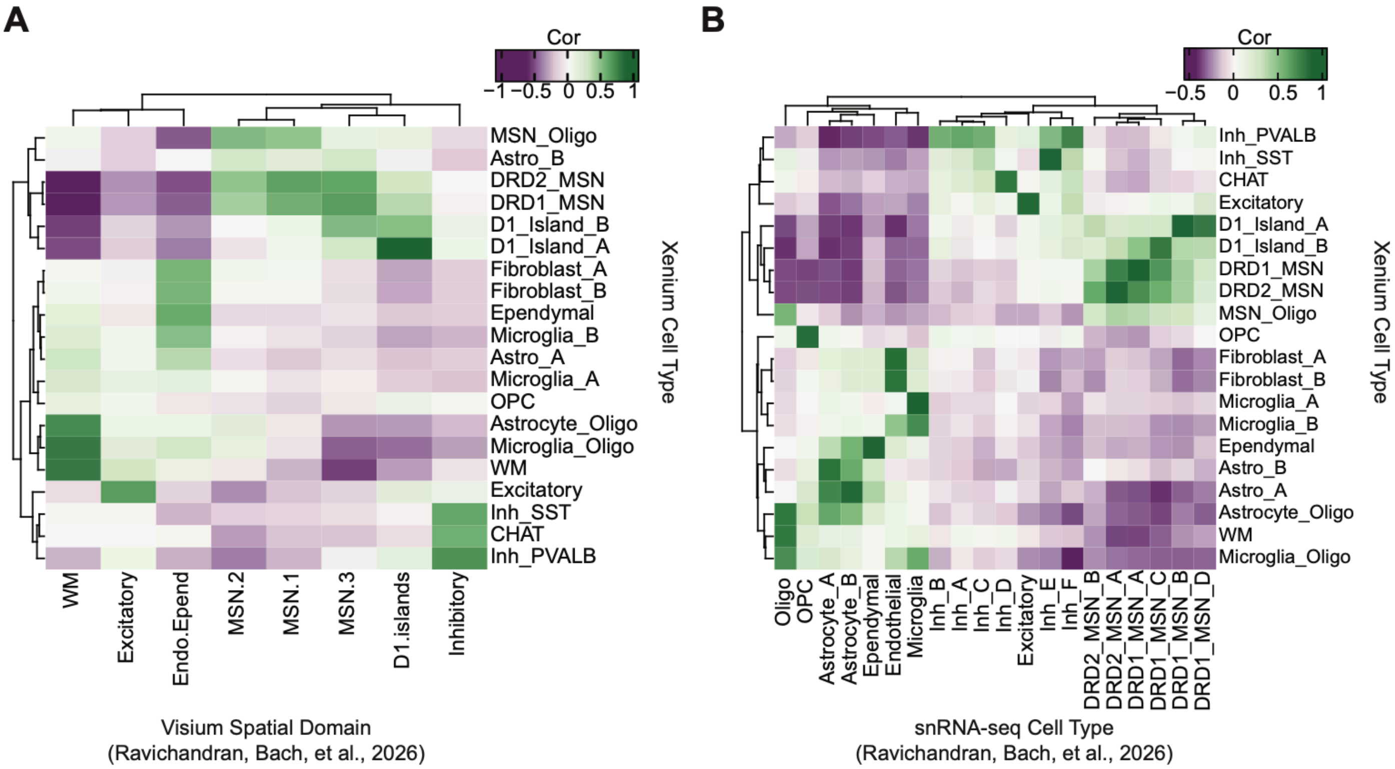

**Figure S5**. **Pairwise alignment and similarity evaluation of consecutive Xenium sections. A.** Initial unaligned placement of the ten consecutive Xenium section pairs from donor Br6660. **B.** Final pairwise placement following *Spateo* alignment. Cells from the two sections in each pair are shown in blue and orange. **C.** Gene-feature similarity to assess the performance between consecutive sections before (blue) and after (green) alignment, evaluated using the median multiscale Pearson correlation coefficient (PCC) and mutual information (MI).

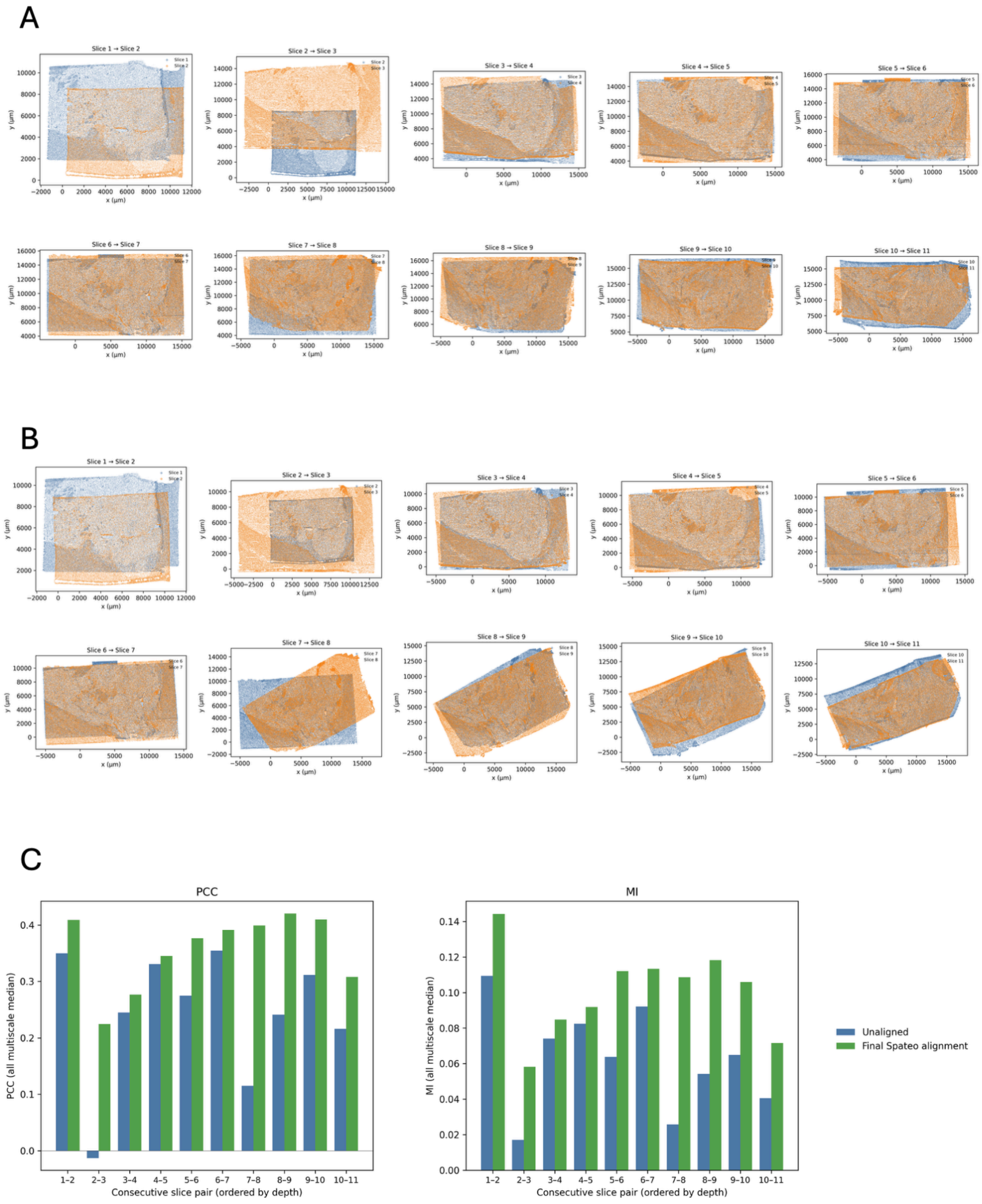

**Figure S6**. **Grid-based analysis of cell type composition across the anterior–posterior axis. A.** A common 5x5 mediolateral–dorsoventral grid overlaid on the 11 aligned Xenium sections from donor Br6660. **B.** Cell type proportions within each of the 25 tiles across the 11 sections, ordered along the AP axis. Each stacked bar represents the cell type composition of one tile region in one section; absent bars indicate that the corresponding tile region did not contain cells in that section. The red box identifies the medial tile regions.

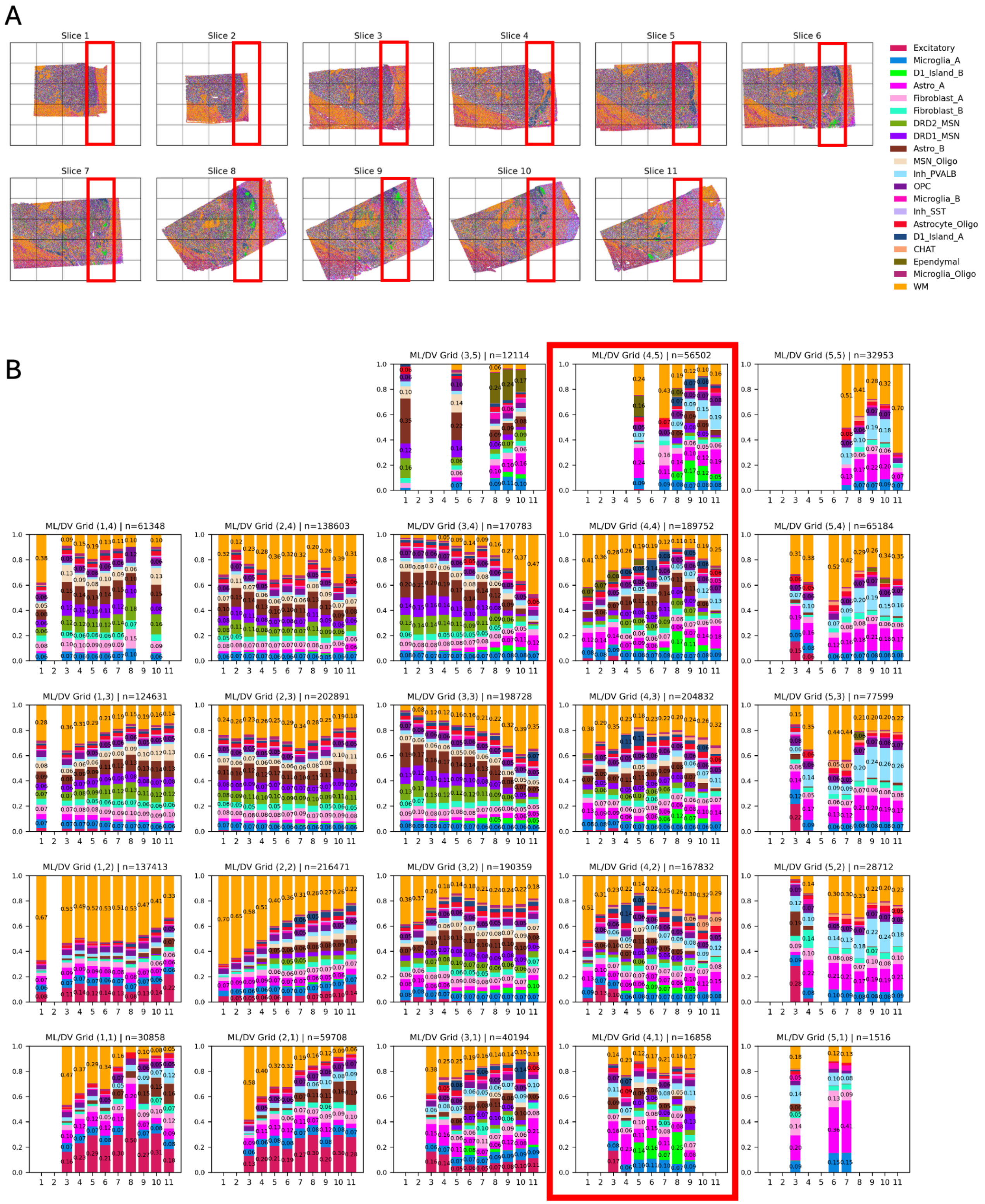

**Figure S7**. **AP expression trends of D1-island marker genes.** Effect-size estimates and 95% confidence intervals are shown for D1-island A and B markers. M0 is the primary model, and M1 is the density-adjusted sensitivity analysis. No displayed marker was significant after Bonferroni correction across 7,320 tests.

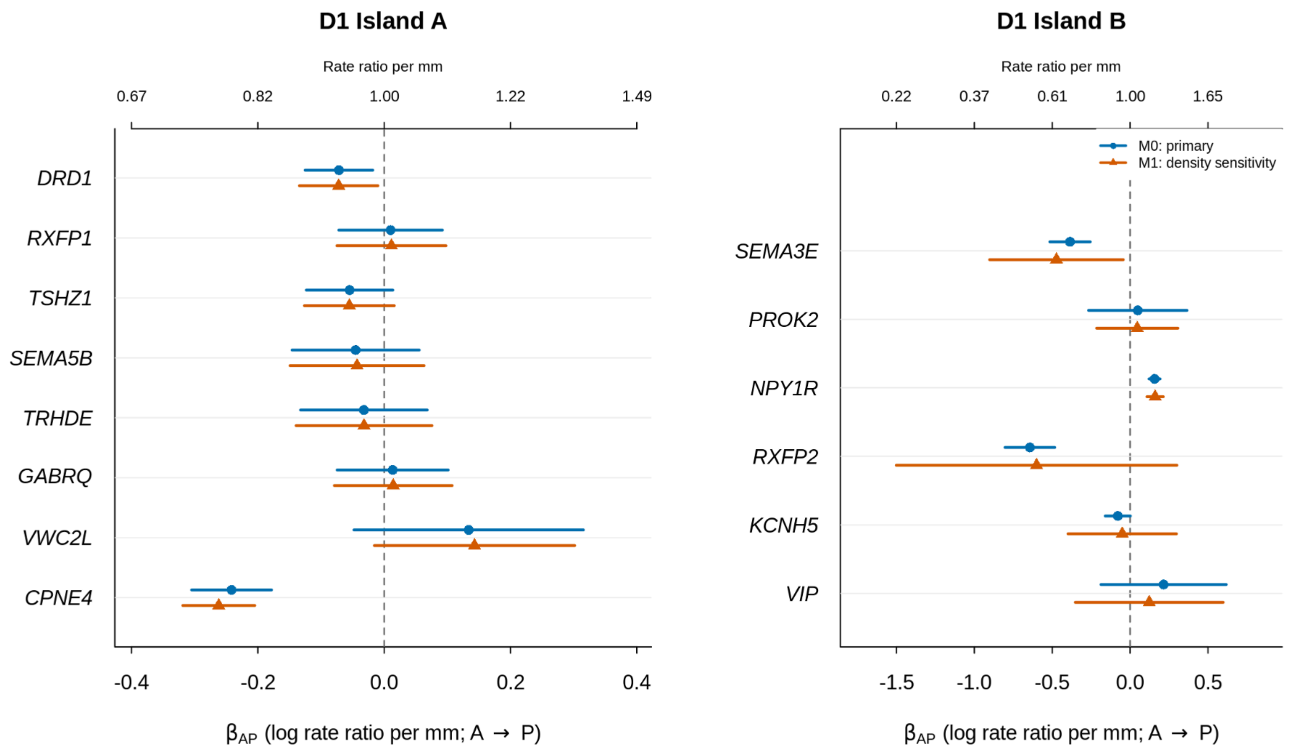

**Figure S8**. Visium HD QC measures. Histograms depicting per-sample distributions of **A**. number of unique molecular identifiers per segmented cell, **B**. number of detected genes per segmented cell, and **C**. percentage of reads mapping to the mitochondrial genome. **D**. Spatial plots colored by cells retained for further analysis or those discarded during QC.

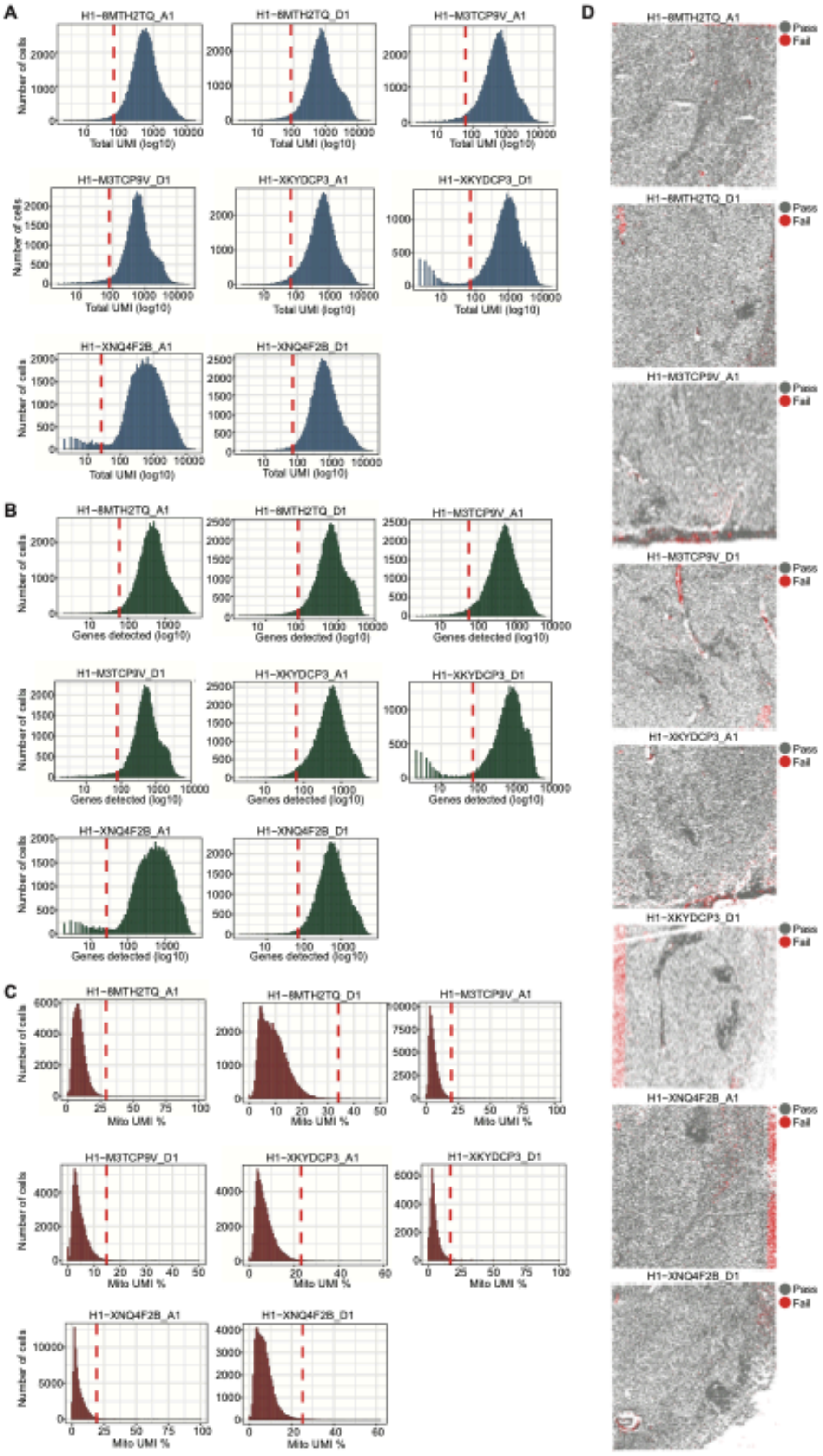

**Figure S9**. **Spatial registration between Visium HD spatial domains and previously reported spatial domains and cell types**. Heatmaps depicting spatial registration results between **A**. Visium HD spatial domains and Visium spatial domains, and **B**. Visium HD spatial domains and snRNA-seq cell types.

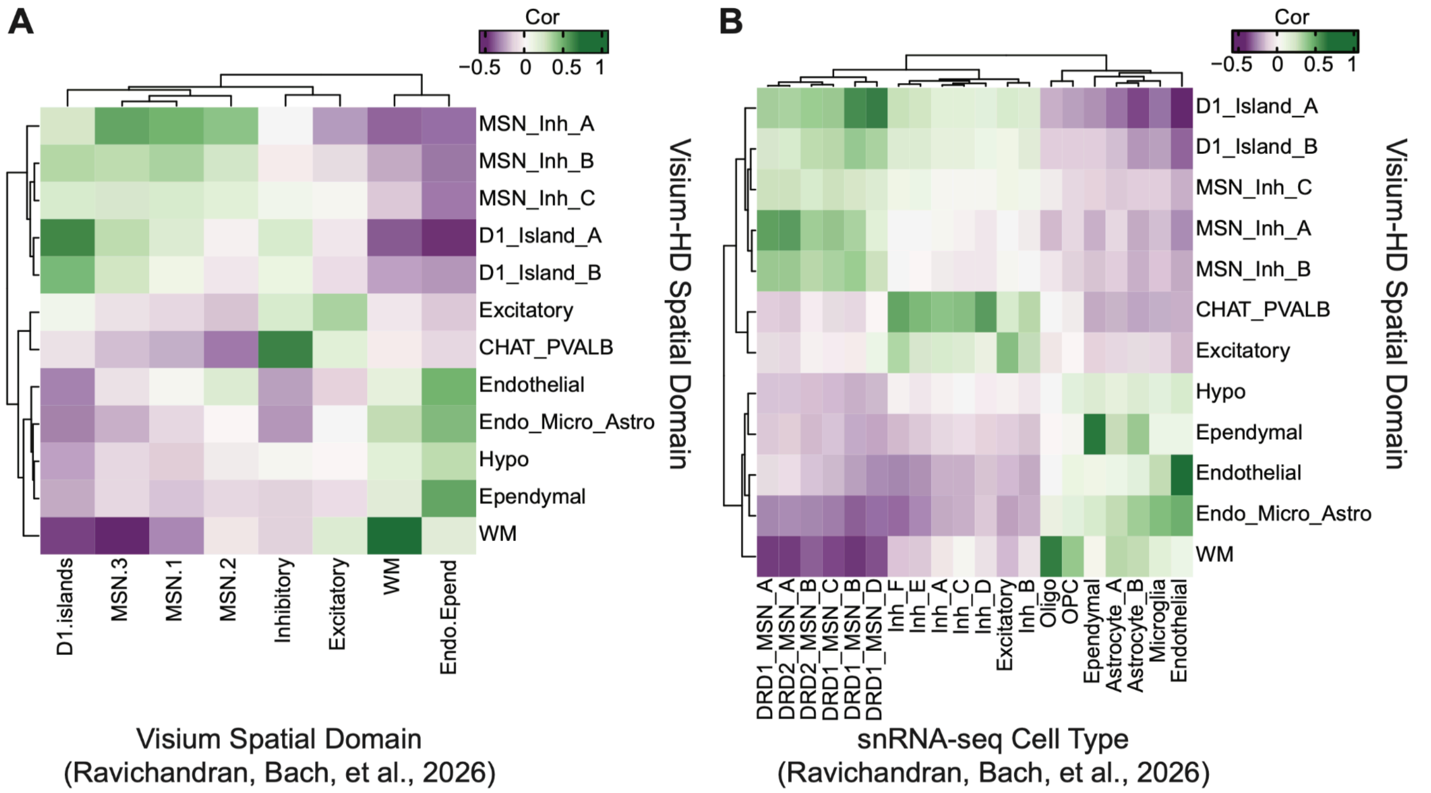

**Figure S10**. Visium HD RCTD label transfer results. **A**. Bargraph showing the percentage of each sample that was labeled as singlet, reject, doublet_certain, or doublet_uncertain by RCTD. **B**. Heatmap depicting the percentage of RCTD first type that is labeled a different cell type (RCTD second type). Only doublets are included in this graph, further demonstrating that cells labeled doublets are likely a single neuronal cell type that happens to include transcripts from nearby non-neuronal cell types.

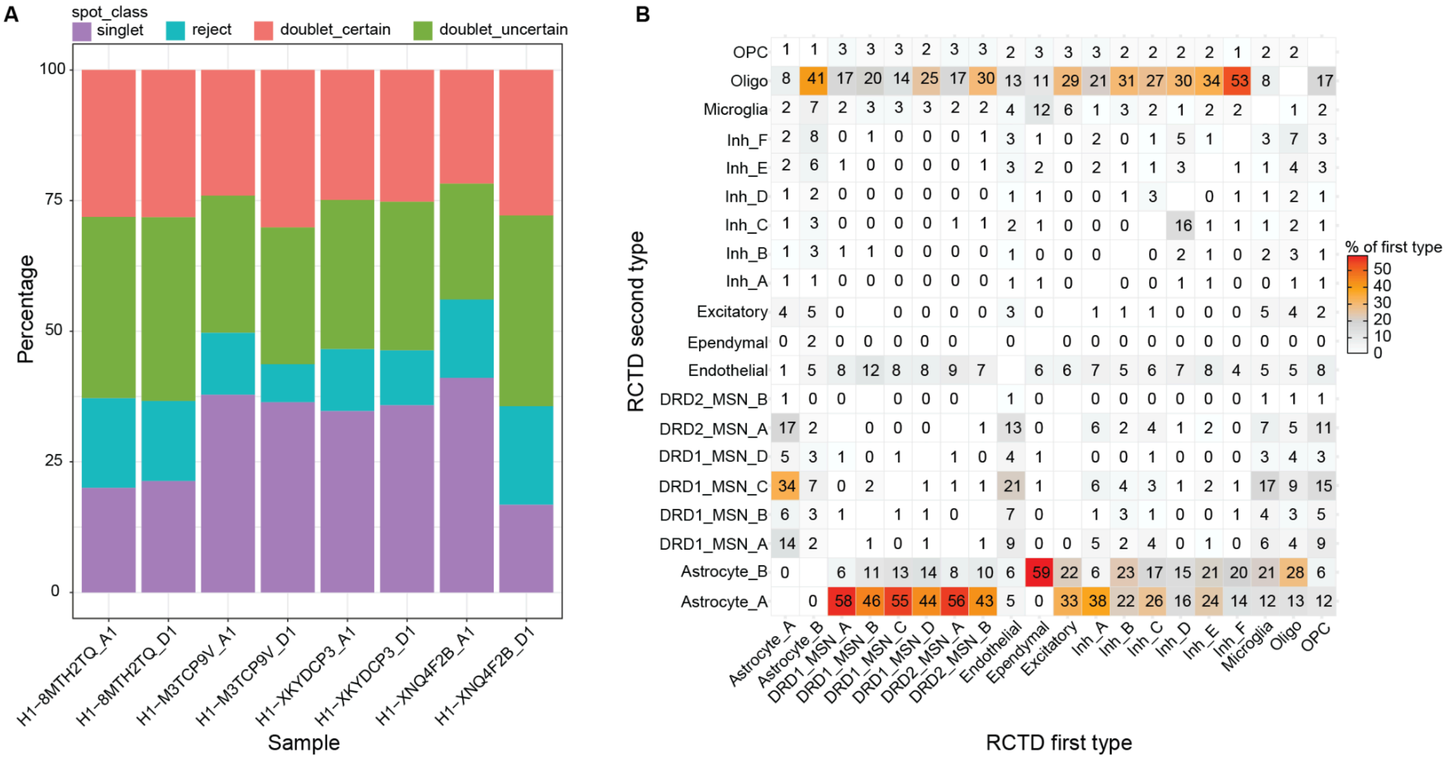

**Figure S11**. **Human NAc snRNA-seq NMF projected into human Visium HD data. A**. Diagram depicting reference and target datasets used in non-negative matrix factorization (NMF) projection. **B**. Spatial plots with Visium HD spatial domains. **C**. NMF loadings for NMF34, specific to DRD1_MSN_B, plotted on Visium HD arrays. **D**. NMF loadings for NMF35, specific to DRD1_MSN_B, plotted on Visium HD arrays. **E**. NMF loadings for NMF44, specific to DRD1_MSN_D, plotted on Visium HD arrays.

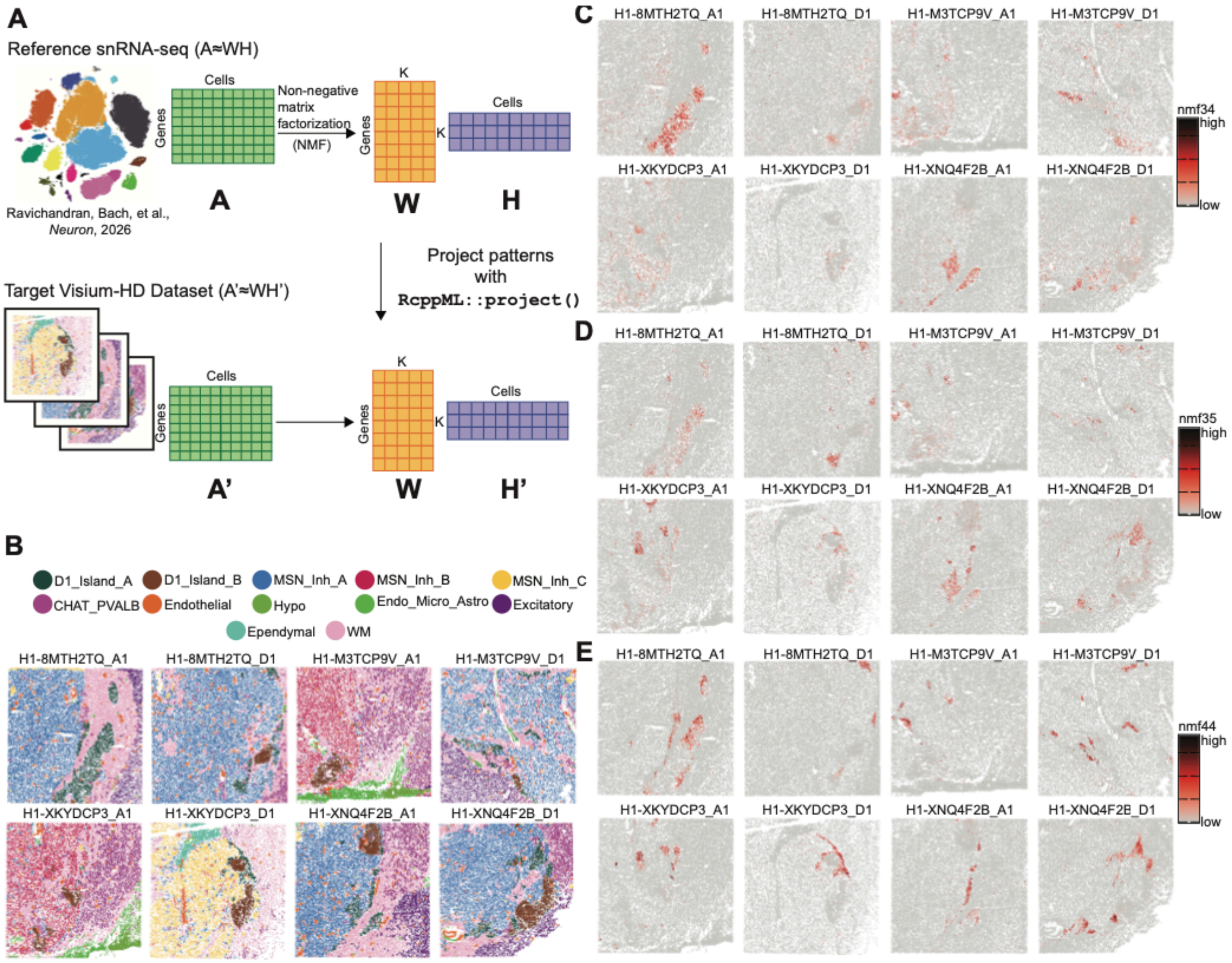

**Figure S12**. RNAscope of D1-island markers. 40x smFISH RNAscope merge of Nac D1_A islands at 2070um level. **A**. High magnifications of DRD1 (magenta), RXFP1 (red) and VIP (yellow) individual channels. **B**. 40x smFISH RNAscope merged of NAc D1_A islands at 3070um level **C**. High magnifications of DRD1 (magenta), RXFP1 (red) and VIP (yellow) individual channels. **D**. 40x smFISH RNAscope merged of NAc D1_B islands at 2090um level. **E**. High magnifications of DRD1 (magenta), PROK2 (red) and SEMA5B (yellow) individual channels. **F**. 40x smFISH RNAscope merge of NAc D1_B islands at 3090um level. **G**. High magnifications of DRD1 (magenta), PROK2 (red) and SEMA5B (yellow) individual channels. **H**. Scale bar 1mm.

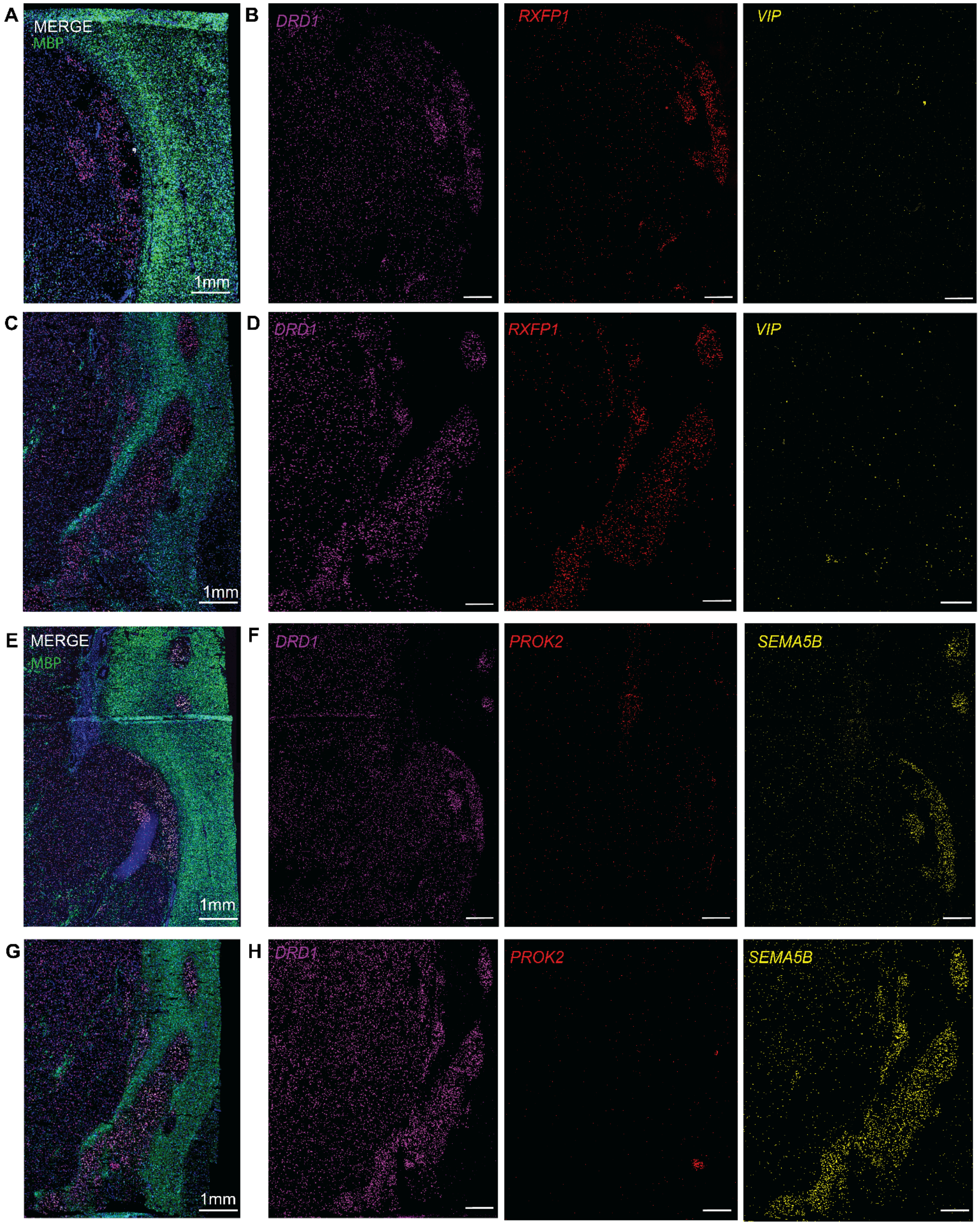

**Figure S13**. **Non-negative matrix factorization of rat snRNA-seq data**[1]. **A**. Cross-validation determines k=53 as optimal rank. **B**. Enrichment of 53 NMF patterns across rat snRNA-seq cell types. We identify NMF10 as a pattern specific to Islands of Calleja and NMF18 as a pattern specific to the Sema5a island subtype. **C**. Normalized NMF loadings for NMF10, a pattern specific to Islands of Calleja, are enriched within Islands of Calleja in a mouse Xenium spatial transcriptomics dataset[1]. **D**. Genes with highest loadings across NMF patterns associated with rat D1-island populations. NMF10 is associated with ICj, NMF18 is associated with Sema5a islands, and NMF11 and NMF24 are associated with Chst9 islands. Sema5a and Chst9 populations are transcriptionally distinct island subtypes in the rat NAc.

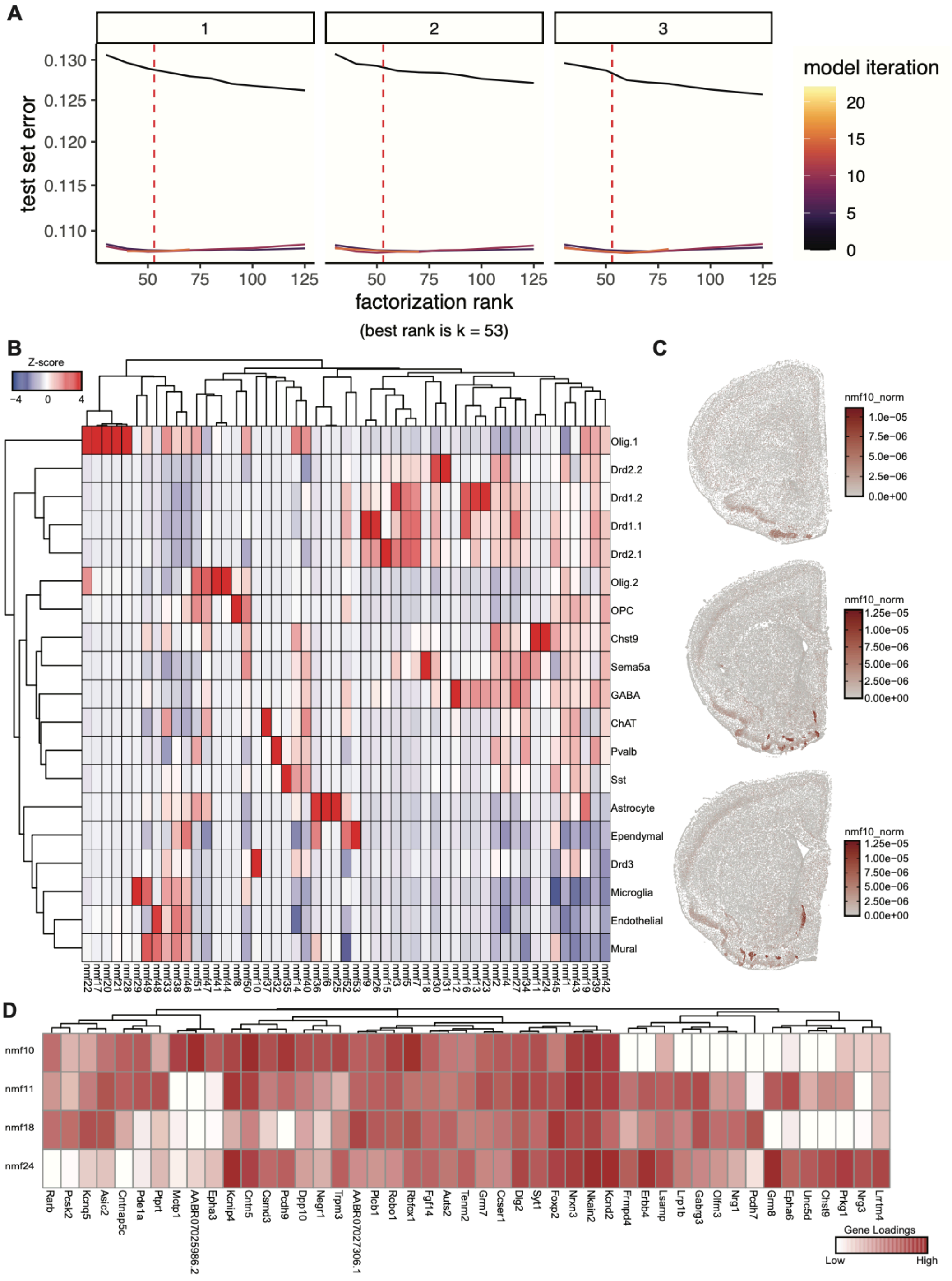

**Figure S14**. **Pairwise alignment of Xenium and Visium HD sections across AP levels.** Each Visium HD section was paired with the nearest Xenium section in AP depth for pairwise *Spateo* alignment. Overlaid spatial coordinates after alignment are shown for **A.** D1_Island_A, **B.** D1_Island_B, and **C.** white-matter cell types. Xenium and Visium HD cells are shown in blue and orange, respectively.

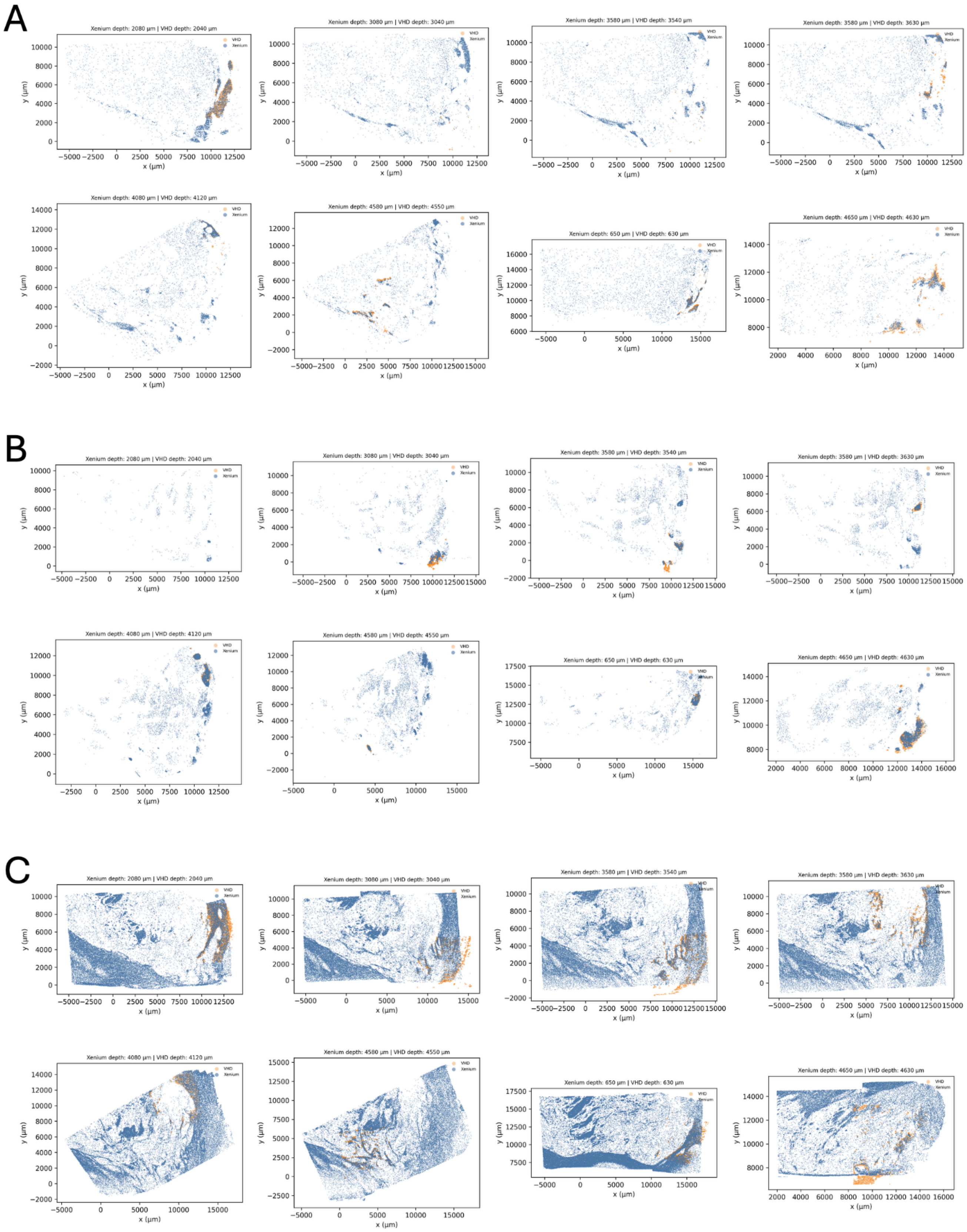

**Figure S15. Spatial heterogeneity of cell type relationships across AP sections and anatomical regions.** **A.** Spatial-relationship summary matrices for all directional cell type pairs in six representative section–region combinations from donor Br6660. The dorsomedial ROI is shown at slice 3, slice 8, and slice 11. Dorsomedial, ventromedial, lateral, and outside-global-ROI regions are shown for slice 8. **B.** Directional spatial relationships from D1_Island_A to D1_Island_B (top) and from D1_Island_B to D1_Island_A (bottom) across AP sections and regions. The corresponding plots show *CRAWDAD* Z-scores across shuffle scales for each AP section and for the lateral, dorsomedial, ventromedial, and outside-global-ROI regions. Horizontal dotted lines indicate the significance threshold (|Z|=3.84).

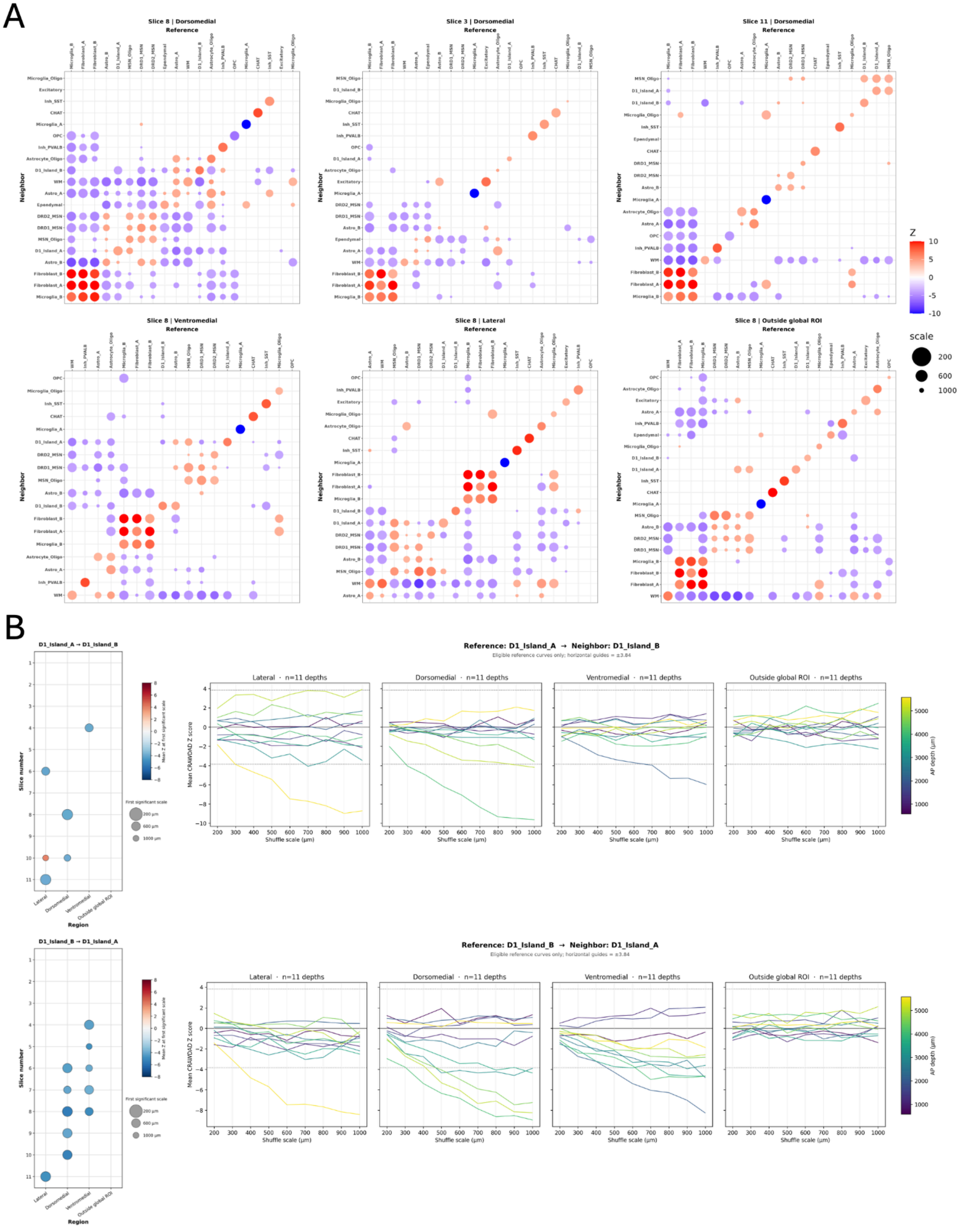
